# Host genetic variation guides hepacivirus clearance, chronicity, and liver fibrosis in mice

**DOI:** 10.1101/2023.03.18.533278

**Authors:** Ariane J. Brown, John J. Won, Raphael Wolfisberg, Ulrik Fahnøe, Nicholas Catanzaro, Ande West, Fernando R. Moreira, Mariana Nogueira Batista, Martin T. Ferris, Colton L. Linnertz, Sarah R. Leist, Cameron Nguyen, Gabriela De la Cruz, Bentley R. Midkiff, Yongjuan Xia, Stephanie A. Montgomery, Eva Billerbeck, Jens Bukh, Troels K.H. Scheel, Charles M. Rice, Timothy P. Sheahan

**Affiliations:** Department of Epidemiology, University of North Carolina at Chapel Hill, Chapel Hill, NC 27599, USA; Copenhagen Hepatitis C Program (CO-HEP), Department of Infectious Diseases, Copenhagen University Hospital, Hvidovre and Department of Immunology and Microbiology, University of Copenhagen, Copenhagen, Denmark; Laboratory of Virology and Infectious Disease, The Rockefeller University, New York, NY, USA; Department of Genetics, University of North Carolina at Chapel Hill, Chapel Hill, NC 27599, USA; Lineberger Comprehensive Cancer Center, University of North Carolina School of Medicine, Chapel Hill, NC 27599, USA; Department of Pathology and Laboratory Medicine, University of North Carolina School of Medicine, Chapel Hill, NC 27599, USA; Department of Medicine, Division of Hepatology, Department of Microbiology and Immunology, Albert Einstein College of Medicine, Bronx, NY, USA

**Keywords:** Animal models, hepatitis c virus, Collaborative Cross, virus-host interactions, viral hepatitis

## Abstract

**Background & Aims:** Human genetic variation is thought to guide the outcome of hepatitis C virus (HCV) infection but model systems within which to dissect these host genetic mechanisms are limited. Norway rat hepacivirus (NrHV), closely related to HCV, causes chronic liver infection in rats but causes acute self-limiting hepatitis in typical strains of laboratory mice, which resolves in two weeks. The Collaborative Cross (CC) is a robust mouse genetics resource comprised of a panel of recombinant inbred strains, which model the complexity of the human genome and provide a system within which to understand diseases driven by complex allelic variation.

**Approach & Results:** We infected a panel of CC strains with NrHV and identified several that failed to clear virus after 4 weeks. Strains displayed an array of virologic phenotypes ranging from delayed clearance (CC046) to chronicity (CC071, CC080) with viremia for at least 10 months. Body weight loss, hepatocyte infection frequency, viral evolution, T-cell recruitment to the liver, liver inflammation and the capacity to develop liver fibrosis varied among infected CC strains.

**Conclusions:** These models recapitulate many aspects of HCV infection in humans and demonstrate that host genetic variation affects a multitude of virus and host phenotypes. These models can be used to better understand the molecular mechanisms that drive hepacivirus clearance and chronicity, the virus and host interactions that promote chronic disease manifestations like liver fibrosis, therapeutic and vaccine performance, and how these factors are affected by host genetic variation.

## Introduction

Hepatitis C virus (HCV) causes chronic infection of the liver in a majority of those infected. If untreated, chronic HCV infection can lead to the development of end-stage liver disease including hepatocellular carcinoma (HCC) and cirrhosis and may require liver transplant^1-4^. Curative treatments, although costly, are available yet undiagnosed cases, access to treatment, reinfection, drug resistance development and a lack of an effective vaccine have complicated efforts for global elimination of HCV as a public health threat^2,5^. In addition, it is not yet clear if cure will prevent the subsequent development of HCC^1,2,4^. While human genetic variation (e.g. *IL28B* locus) is thought to guide both the kinetics of spontaneous clearance and chronic infection, animal model systems within which to dissect these host genetic mechanisms are not available^6^. Thus, our understanding of how host genetic variation affects HCV pathogenesis, the development of chronicity, immunity and vaccine efficacy is hampered by the lack of robust immune competent small animal models.

HCV-related hepaciviruses have been discovered in multiple animal species including horses, non-human primates, wild rodents and bats^7^. In 2014, Norway rat hepacivirus (NrHV) was discovered in rats in New York City to cause chronic liver infection^8^. Many basic aspects of NrHV biology are shared with HCV, including hepatotropism, genome organization, dependence on a liver-specific micro RNA (miR-122) for replication and the use of scavenger receptor B-1 for entry^9-11^. In common lab strains of mice, NrHV causes subclinical self-limiting hepatotropic infection that is cleared in 14-21 days but chronic infection can be initiated via transient depletion of CD4 T-cells prior to infection^9^. Models of chronic hepacivirus infection in fully immune competent mice without immune manipulation are not yet available.

The Collaborative Cross (CC) is a robust mouse genetics resource created by the systematic breeding of eight founder strains encompassing classic inbred, disease model and wild-derived models resulting in the generation of a panel of recombinant inbred (RI) strains, which together model the complexity of the human genome and provide a system within which to understand disease driven by complex allelic variation^12^. The CC contains greater than 90% of the common genetic variation in mouse resources through the presence of uniformly distributed SNPs (>40 million) and indels (>4 million) across the various genomes^13,14^. When applied to virology, the CC has been a powerful tool to both identify better models of human disease, as shown with Ebola virus^15^, severe acute respiratory syndrome associated coronavirus (SARS-CoV)^16^, West Nile virus^17,18^, SARS-CoV-2^19^, and influenza A virus^20^, and also to map the genetic determinants responsible for complex phenotypic traits^13^.

Here, we describe multiple CC mouse models of chronic hepacivirus pathogenesis. We show that host genetic variation affects hepacivirus chronicity, the host response to infection, hepatocyte infection frequency, liver inflammation, liver fibrosis and viral evolution thus recapitulating many aspects of HCV disease biology. This system can be used to better understand the host genes and molecular mechanisms that guide hepacivirus pathogenesis, liver fibrosis, therapeutic and vaccine performance.

## Results

### Host genetics is a determinant of acute hepacivirus clearance

Given the profound impact human genetic variation can have on HCV infection, we aimed to determine if host genetic variation would affect NrHV pathogenesis and infection kinetics in mice (Fig. 1A). We infected ten different CC mouse strains with a mixture of two sub-populations of mouse derived NrHV inoculum, NrHV-A and -B along with a putative mouse adapted NrHV-B^9^ (B_SLIS_) at a ratio of 1:1:1 (A:B:B_SLIS_) (Fig. S1). We utilized this mixture to give the best possible foundation for persistency not knowing which genotype would be optimal in the CC. After infection, we monitored viremia for 4 weeks which is 1-2 weeks after WT C57BL/6J (WT) mice typically clear. All strains of mice became infected following NrHV inoculation (Fig. S2), but most CC strains and WT had cleared the infection by week 4. In contrast, CC040, CC046, CC071 and CC080 remained viremic at week 4 (Fig. 1B) and both CC071 (P = 0.02) and CC080 (P = 0.002) had significancy elevated levels of viremia. Concurrently, CC071 (P = <0.0001) and CC080 (P = 0.001) intrahepatic viral RNA was significantly elevated over WT (Fig. 1C). Gene expression of Mx1, a typical interferon stimulated gene (ISG), was elevated in the livers of CC strains that remained viremic, a sign that sustained virus replication continued to instigate the host (Fig. 1D). All together, these data demonstrate that host genetic variation influences acute hepacivirus clearance.

**Figure 1.**
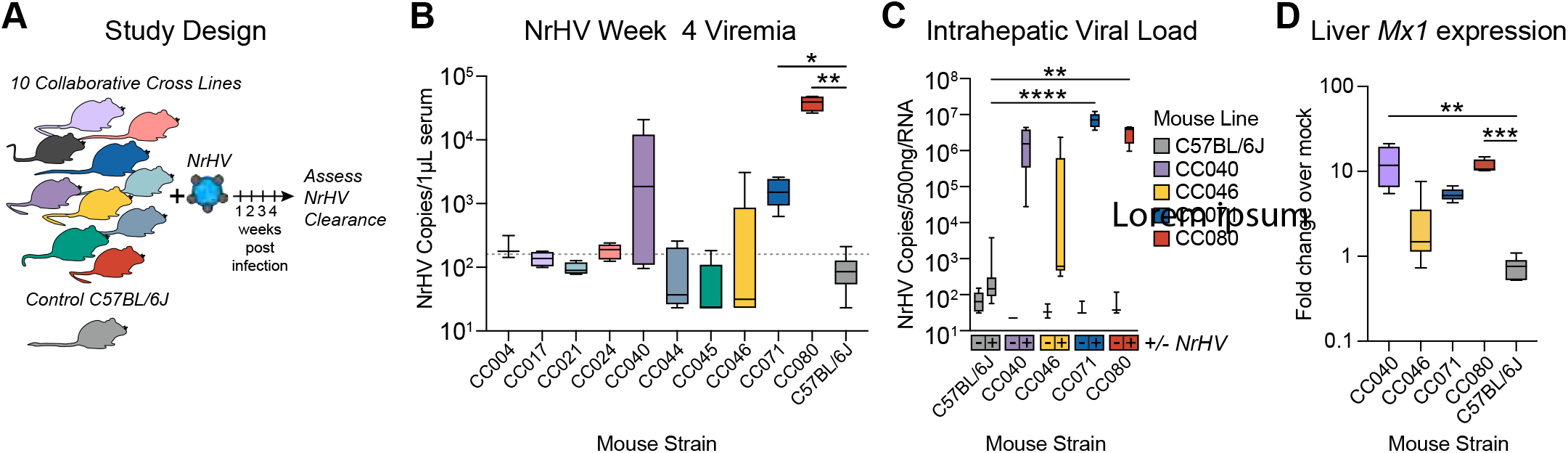
Host genetics is a determinant of acute hepacivirus clearance. **(A)** Study design. Ten Collaborative Cross (CC) strains and control C57BL/6J female mice 9-13 weeks in age were infected with 1×10^5^ genome equivalents of recombinant NrHV or negative control PBS via retroorbital injection and bled weekly to monitor viremia. Mouse numbers per strain were: C57BL/6J N = 8, CC004 N = 4, CC017 N = 4, CC021 N = 4, CC024 N = 5, CC040 N = 5, CC044 N = 5, CC045 N = 5, CC046 N = 6. CC071 N = 5, CC080 N = 4. For all strains PBS mock infected N = 3. **(B)** Week 4 viremia as determined by qRT-PCR of viral RNA isolated from serum. The dotted line indicates the limit of quantitation. **(C)** Intrahepatic viral load 4 weeks post infection by qRT-PCR using 500 ng total liver RNA. Numbers of mice per group: C57BL/6J (8 NrHV, 4 Mock), CC040 (4 NrHV, 3 mock), CC046 (6 NrHV, 3 mock), CC071 (5 infected, 3 mock), CC080 (4 infected, 3 mock). **(D)** *Mx1* intrahepatic gene expression 4 weeks by qRT-PCR using 500ng total RNA. Data is expressed as fold change over mock infected by ΔΔCT method. For B and D, asterisks indicate statistical significance as determined by Kruskal-Wallis test. For C, asterisks indicate statistical significance by Two-Way ANOVA with a Dunnett’s multiple comparison test.

### Host genetics is a determinant of hepacivirus chronicity

We then further characterized a subset of CC strains with sustained NrHV replication at 4 weeks. CC046, CC071 and CC080 are genetically distinct with notably different genomic architectures including at loci likely to be important for the antiviral response such as major histocompatibility complex (MHC), T-cell activation, and type I and III interferons (IFN) (Fig. S3). We infected WT, CC046, CC071 and CC080 mouse strains with NrHV as done above and monitored them for 10 months (Fig. 2A). Interestingly, body weight loss was only observed in CC071 infected mice and only during the first week (Fig. S4 and S5). As expected, WT cleared virus within 3 weeks (Fig. 2B). In contrast, CC046 had sustained replication during weeks 1-4 but ultimately all mice cleared virus by week 7 (Fig. 2B). Notably, the majority of CC071 mice became chronically infected with quantifiable viremia until the termination of the study at week 38-39 (Fig. 2B). CC080 mice had a variety of virologic outcomes where some mice cleared virus in middle (4-7 weeks) or late (8-22 weeks) times but only a minority became chronically infected (Fig. 2B). During the first two weeks of infection, the levels of viremia in CC071 (week 1 P = 0.007, week 2 P = 0.001) and CC080 (week 1 P = 0.036, week 2 P = 0.02) were significantly higher than WT and by week 3, viremia in all three CC strains exceeded that in WT mice (CC046 P = 0.006, CC071 P = 0.007, CC080 P = 0.04) (Fig. 2C). While all CC strains had measurable viremia in week 5, only CC071 (P = 0.01) remained significantly elevated over WT (Fig. 2C).

**Figure 2.**
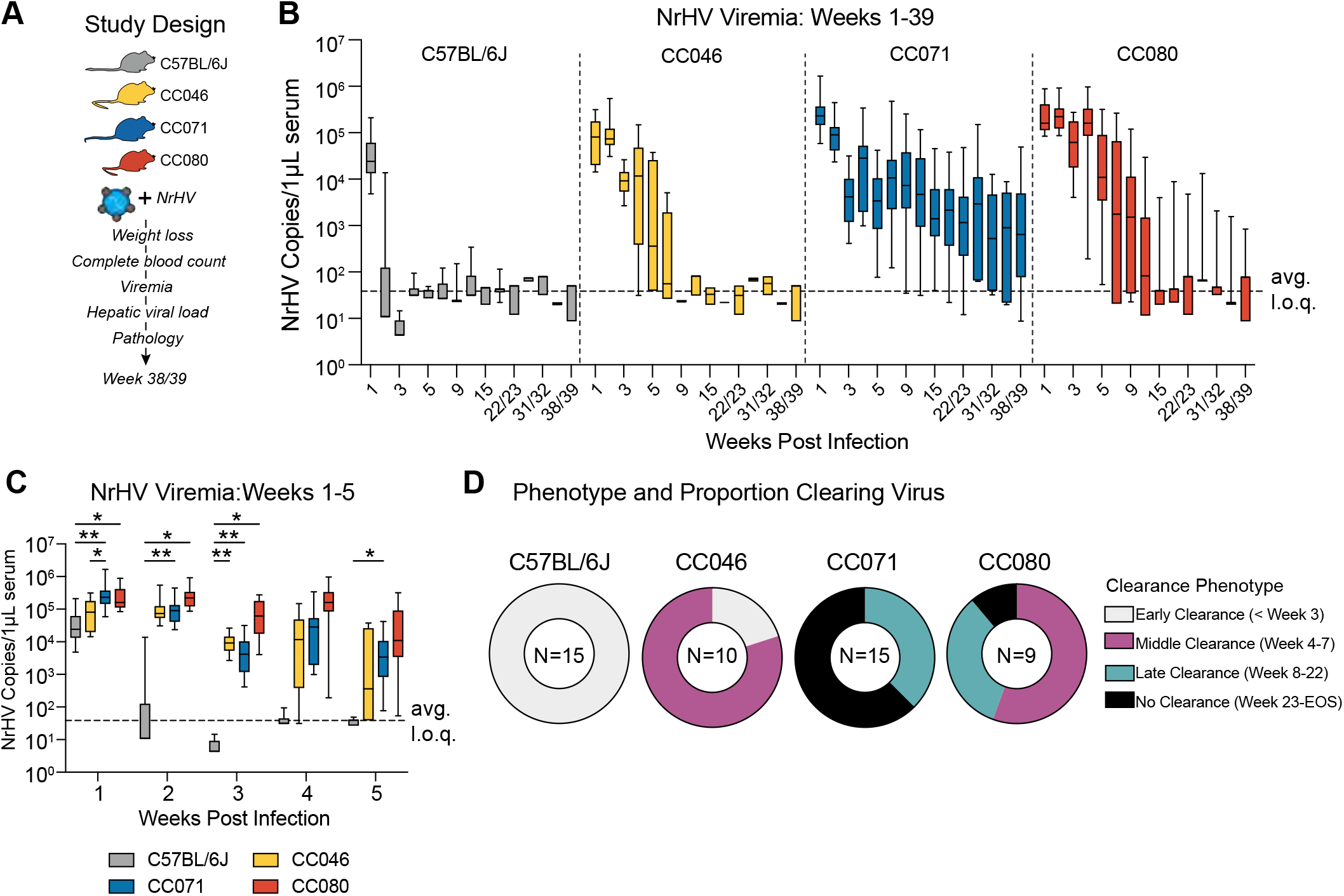
Host genetics is a determinant of hepacivirus chronicity. **(A)** Study Design. C57BL/6J, CC046, CC071 and CC080 mice were infected with 1×10^5^ genome equivalents of recombinant NrHV or negative control PBS via retroorbital injection and viremia was monitored weekly or monthly. **(B)** NrHV viremia for weeks 1-39 determined by qRT-PCR of RNA from serum. See Table S2 for the numbers of animals per strain per time. **(C)** NrHV Viremia for weeks 1-5. Levels of viral RNA in serum were measured by qRT-PCR. C57BL/6J Time (N): week 1 (19), week 2 (15), week 3 (15), week 4 (15), week 5 (15). CC046 Time (N): week 1 (13), week 2 (10), week 3 (10), week 4 (10), week 5 (10). CC071 Time (N): week 1 (22), week 2 (20), week 3 (18), week 4 (19), week 5 (19). CC080 Time (N): week 1 (13), week 2 (10), week 3 (10), week 4 (10), week 5 (10). Asterisks indicate statistically significant differences by Two-Way ANOVA Tukey’s multiple comparisons test. **(D)** Proportion of mice per strain that clear virus early (< 3 weeks), middle (week 4-7), late (week 8-22) or do not clear (viremic week 23 to end of study). Only mice that survived to the end of the study were considered for this analysis. For B and C, the dotted line indicates the average limit of quantitation. All data for B-D, was compiled from two independent experiments.

To assess infectivity in serum, we passively transferred serum from 10-weeks post infection to naïve WT animals and then monitored viremia over time (Fig. S6). As expected, transfer of serum from aviremic mice (WT and CC046) did not transfer infection to recipient animals (Fig. S6B, S6C) but transfer of serum from viremic CC071 (Fig. S6D) and CC080 (Fig. S6E) mice resulted in infection in recipient animals. Thus, viral RNA associated with significant viremia is infectious. Interestingly, when examining viremia per animal, chronically infected animals experienced periods of relatively consistent levels of viremia followed by drastic increases or decreases suggesting an interplay among virus and the host response (Fig. S7). Altogether, by varying host genetics, a spectrum of virologic outcomes were observed following NrHV infection where all WT mice clear early (< 3 weeks), most CC046 mice clear between weeks 4-7, the majority of CC071 became chronically infected, and CC080 displayed phenotypes seen in both CC046 and CC071(Fig. 2D).

### Host genetics determines infection frequency and viral dynamics in the liver

To complement our virologic measures in peripheral blood, we next extended our studies to the liver. Labeling viral genomic RNA via *in situ* hybridization (ISH) revealed a variety of phenotypes. First, clearance phenotypes observed in the periphery were similarly observed in WT and CC046 and most CC071 and some CC080 remained viral RNA ISH positive for the duration of the study (Fig. 3A). Second, the amount of viral RNA per cell varied widely especially at 2dpi where the amount of viral RNA per cell was greatest in CC071 (Fig. 3A). Third, the average infection frequency varied greatly among the different strains. On 2dpi, the infection frequency for CC046 was 22%, more than 10-fold greater than other strains (Fig. 3B). By 7dpi, the infection frequency in CC071 mice had increased similar levels, still significantly greater than both WT and CC080 mice. By 28dpi, the infection frequency in CC071 mice had increased even further (average frequency = 38.1%) but decreased to levels seen at earlier times by the end of the study (Fig. 3B). In paired tissue samples, we quantitated viral RNA via qRT-PCR. As seen with ISH, CC046 had significantly elevated levels of NrHV RNA at 2dpi (P = 0.03-0.04) compared to other CC strains (Fig. 3C). Like the viremia and ISH data, CC071 had elevated levels of intrahepatic viral RNA over WT and CC046 at both 28dpi and 272dpi. Altogether, the intrahepatic virologic data was concordant with that observed in the periphery but also revealed new insights into the impact of host genetic variation on infection frequency and replication dynamics.

**Figure 3.**
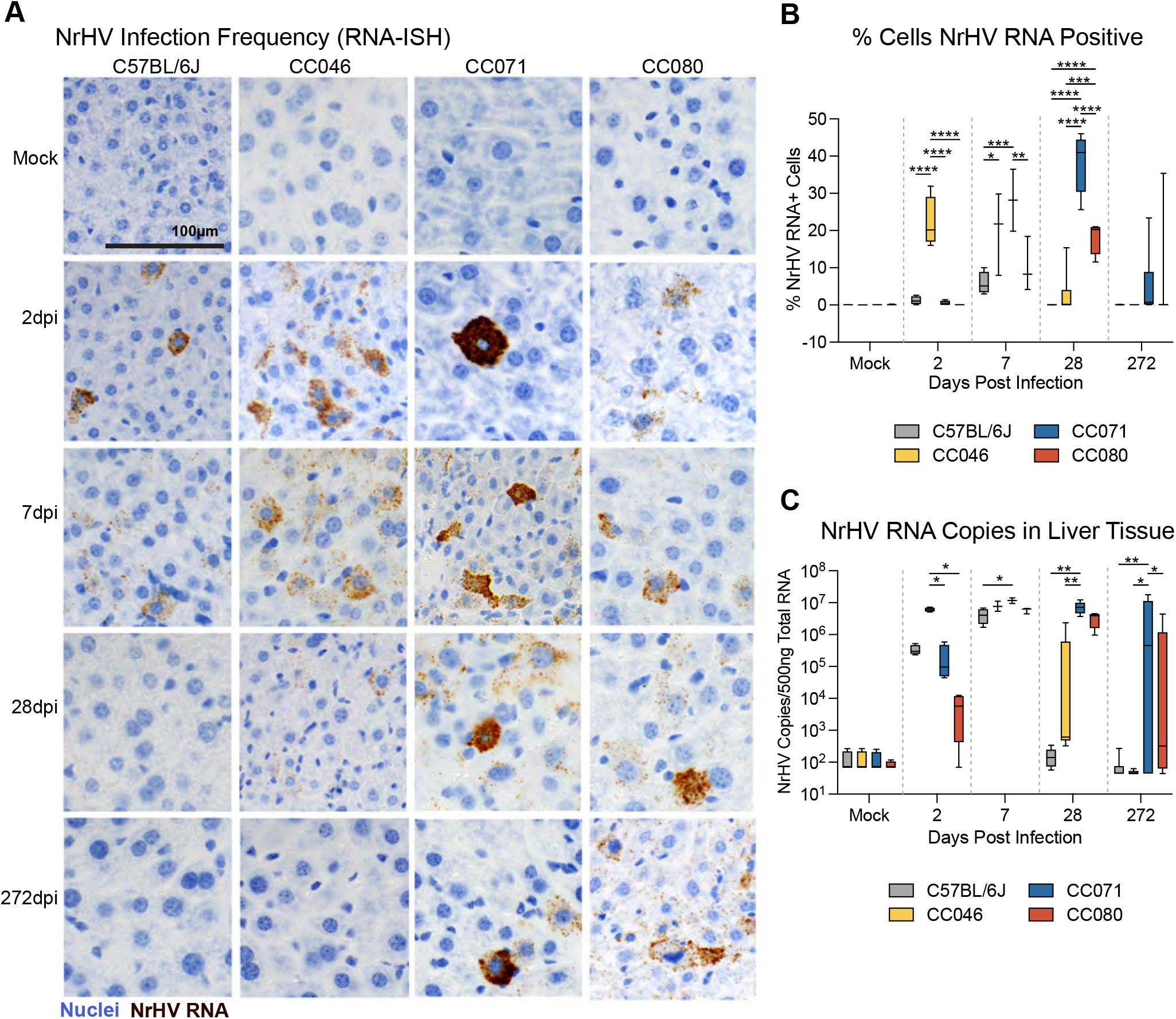
Host genetics determines infection frequency and viral dynamics in the liver. **(A)** NrHV viral RNA *in-situ* hybridization in liver tissue sections from mock or NrHV infected C57BL/6J, CC046, CC071 or CC080 mice 2, 7, 28 or 272 dpi. Nuclei are stained blue and viral RNA is labeled brown. **(B)** Percent cells NrHV RNA positive via RNAscope *in situ* hybridization quantified using Definiens Architect. C57BL/6J Time (N): Mock (6), 2dpi (4), 7dpi (4), 28dpi (5), 272dpi (7). CC046 Time (N): Mock (7), 2dpi (4), 7dpi (3), 28dpi (6), 272dpi (5). CC071 Time (N): Mock (7), 2dpi (4), 7dpi (2), 28dpi (5), 272dpi (15). CC080 Time (N): Mock (6), 2dpi (4), 7dpi (3), 28dpi (4), 272dpi (8). **(C)** NrHV genome copy number in 500ng total liver RNA by qRT-PCR. C57BL/6J Time (N): Mock (4), 2dpi (4), 7dpi (4), 28dpi (5), 272dpi (7). CC046 Time (N): Mock (4), 2dpi (4), 7dpi (3), 28dpi (6), 272dpi (5). CC071 Time (N): Mock (4), 2dpi (4), 7dpi (2), 28dpi (5), 272dpi (15). CC080 Time (N): Mock (4), 2dpi (4), 7dpi (3), 28dpi (4), 272dpi (8). Asterisks in B and C indicate statistical significance by Two-Way ANOVA Tukey’s multiple comparisons test.

### Immune dysregulation is associated with chronic hepacivirus infection

Given the disparate outcomes we had observed thus far, we next aimed to determine if host genetics differentially affected immune responses. First, we performed completed blood count on longitudinal peripheral blood samples from the studies described above. The numbers of total lymphocytes were largely unaffected over time comparing infected and strain matched mock mice (Fig. S8A and B). In contrast, neutrophils were significantly and consistently elevated in NrHV infected CC071 compared to strain matched mock mice in weeks 1 and 2 (Fig. S8C and D). To better understand intrahepatic responses, we labeled liver tissue sections for CD4 positive cells since CD4 T-cells were previously shown to be important for clearance in WT mice^9^. On 2dpi, clusters of CD4+ cells were observed in the parenchyma and on portal tracts in WT mice, who most rapidly clear virus, whereas the labeling pattern remained unchanged from mock for all infected CC strains (Fig. 4). By 7dpi, periportal CD4+ cells were prominent in WT, CC046 and CC071 mice yet the staining pattern for CC080 remained similar to that in uninfected. By 28dpi, clusters of CD4+ cells in periportal and parenchymal space were readily apparent in all infected CC strains, yet the pattern in WT mice, who had all cleared virus by this time, were like mock. By 39 weeks post infection, WT mice remained unremarkable, but clusters of CD4+ cells around portal tracts remained in all chronically infected CC strains and even in CC046 that had cleared after week 7.

**Figure 4.**
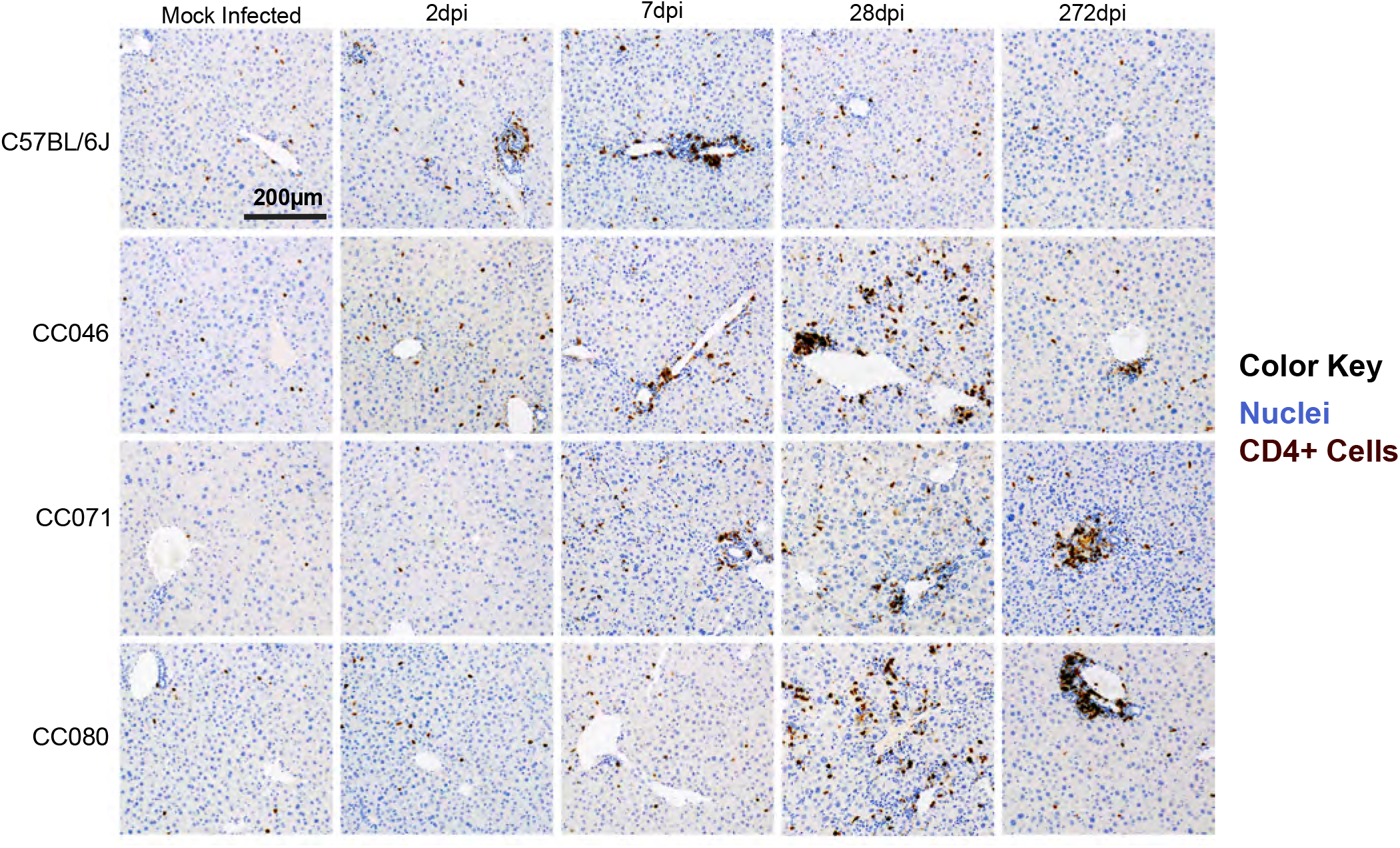
CD4 cell recruitment is delayed in CC mouse strains. CD4 cell labeling in liver tissue sections from mock and NrHV infected animals at 2, 7, 28, and 272dpi with CD4 cells (brown) and nuclei (blue). Representative images of mouse strains and timepoints are shown.

### Dysregulation of the transcriptome in strains susceptible to prolonged infection

To determine if early defects in the host response were associated with chronic infection, we analyzed the transcriptional responses (Fig. 5) in the liver of WT and CC mouse strains on 7dpi when significant virologic and immunologic differences were observed. Importantly, via principal component analysis (PCA), we confirmed the global responses among mock and infected groups differed and like samples tended to cluster together (Fig. 5A). The numbers of significantly regulated genes varied with host genetics with CC071 regulating more genes than all other strains (Fig. 5B). Pathway analysis of significantly regulated genes (<1.5 Log fold change over mock, P = 0.05) revealed the differential regulation of canonical pathways associated with innate and adaptive immunity among WT and CC strains (Fig. 5C). In Figure 5D, we show the expression fold change data for select canonical pathways including Pathogen Induced Cytokine Storm Signaling, LXR/IL-1 Mediated Inhibition of RXR Function, Th1 Signaling and LXR/RXR Activation. For all affected pathways shown, the gene expression patterns for CC071 are quite different than those in WT and even those of other CC strains with a noted lack of expression in innate immune (e.g. Tlr1, 6, 7, and 13), chemokine (e.g. Ccl6, Cxcl1, Cxcl10), adaptive immune (e.g. *Cd247*, *Cd3*) and Il-1/inflammasome (e.g. Il1b, Il1rn, IL18rap, NLRP3, etc.) related genes. Coupled with the CD4+ cell labeling data above, collectively these data show that CC strains with prolonged and chronic infection are dysregulated in key innate and adaptive immune pathways as compared to WT mice which most rapidly clear virus.

**Figure 5.**
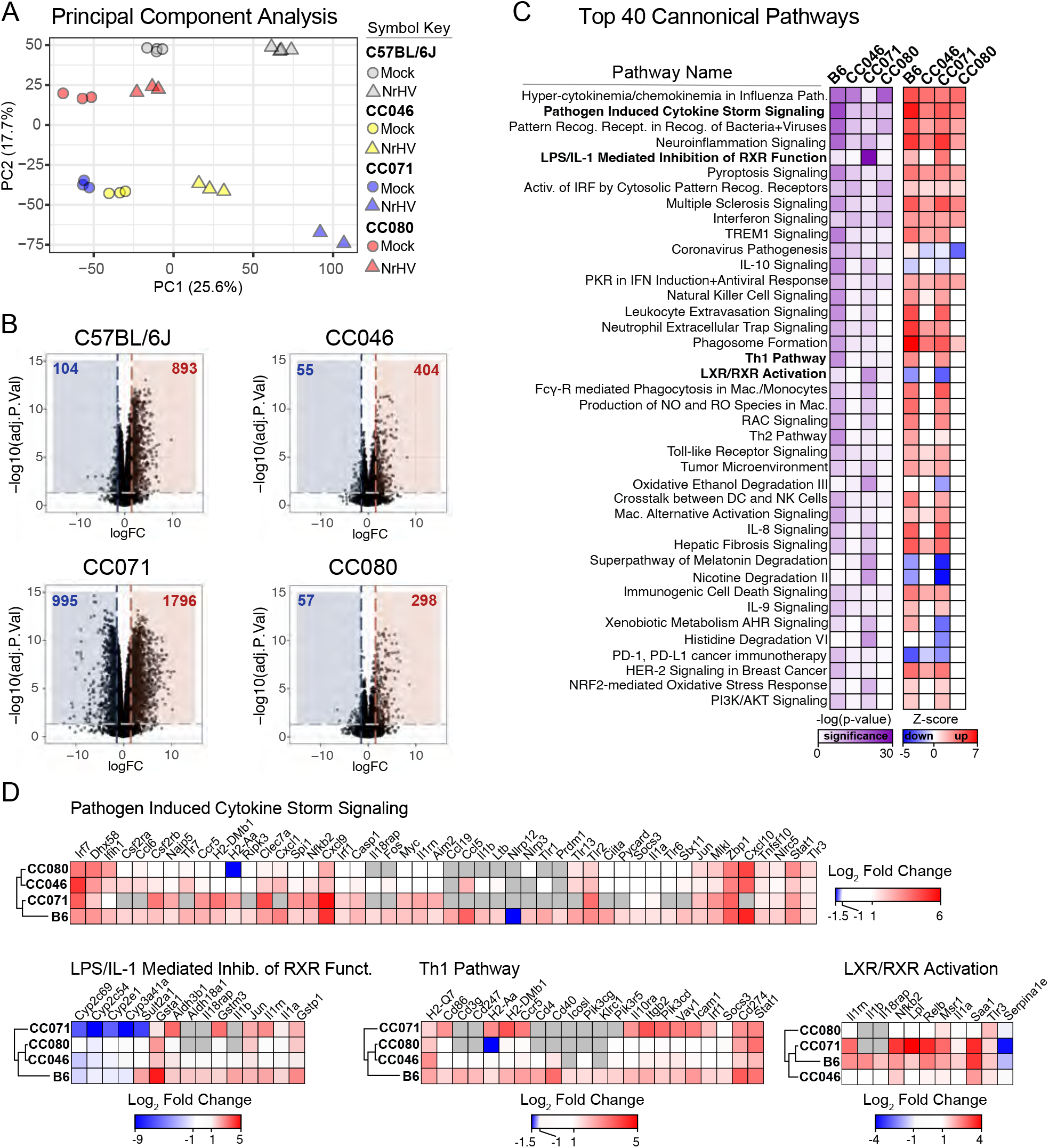
Dysregulation of the transcriptome in strains susceptible to chronic infection. Total RNA from mock and NrHV infected liver tissue was deep sequenced by Illumina HiSeq 4000. For all groups, N = 3 except for CC071 NrHV infected N =2. **(A)** Principal component analysis of expression data for mock and infected groups. **(B)** Top 40 Regulated Canonical Pathways by Ingenuity IPA. The significance (-log(p-value) and predicted regulation (Z-score) for each pathway is indicated by the scale bar. **(C)** Expression of significantly regulated genes (>1.5 Log fold change P < 0.05) over mock in select Canonical pathways. Heat maps were generated in Morpheus.

### Viral evolution is driven by host genetics

To determine if viral evolutionary patterns varied with host genetics, we deep sequenced the complete viral open reading frame (ORF) in longitudinal serum samples from select mice that either cleared virus or developed chronic infection (Fig. S9). Virus populations from chronically infected animals (CC071 #41, 43, 46; CC080 #58, 63) evolved over time, as evidenced by the continued animal specific branching and increase in branch length in the dendrogram (Fig. S10A). In contrast, virus populations from CC046 who cleared virus the earliest among CC strains evolved the least (Fig. S10A). Proteins/regions of the genome under positive selection accumulate non-synonymous changes at increased frequency compared to synonymous changes. Thus, we then compared the non-synonymous and synonymous changes (dN/dS) per viral ORF. Regardless of host genetic background and unlike other portions of the genome, the dN/dS ratio for structural proteins E1 and E2, as well as p7 was increased at all times suggestive of positive selection (Fig. S10B).

We next analyzed the mutational patterns per mouse (Fig. 6). CC071 infected mice (Fig 6A), had more nonsynonymous changes compared to viruses from CC080 (Fig. 6B) or CC046 (Fig. 6C) animals no matter infection outcome. In all three CC mouse strains, the mouse adapted NrHV-B_SLIS_ variant was rapidly selected for likely due to its mutations T190S, V353L, F369I and N550S, which appear to adapt the virus to WT mice^9^. In general, viruses from mice that resolved infection had the fewest additional substitutions but several mutations residing in an MHC-I epitope conserved among mice and rats (T184A, S191F, V196A) were observed in viruses from all strains^21^. In addition, either A698V or C715S appeared to be selected in p7 but rarely together. Interestingly, N550S, predicted to disrupt a glycosylation site, reverted in two of three chronically infected CC071 mice concomitantly with the disruption of another predicted glycosylation site at residues 504-506. Substitution of residue 500 (3/3) and 570 (2/3) was also observed in chronically infected CC071 mice. Further, unique changes linked to persistence were observed in CC071 (R984H in NS3) and CC080 (P1446S/L). Thus, viral evolution, clearance and chronicity were closely linked to host genetics.

**Figure 6.**
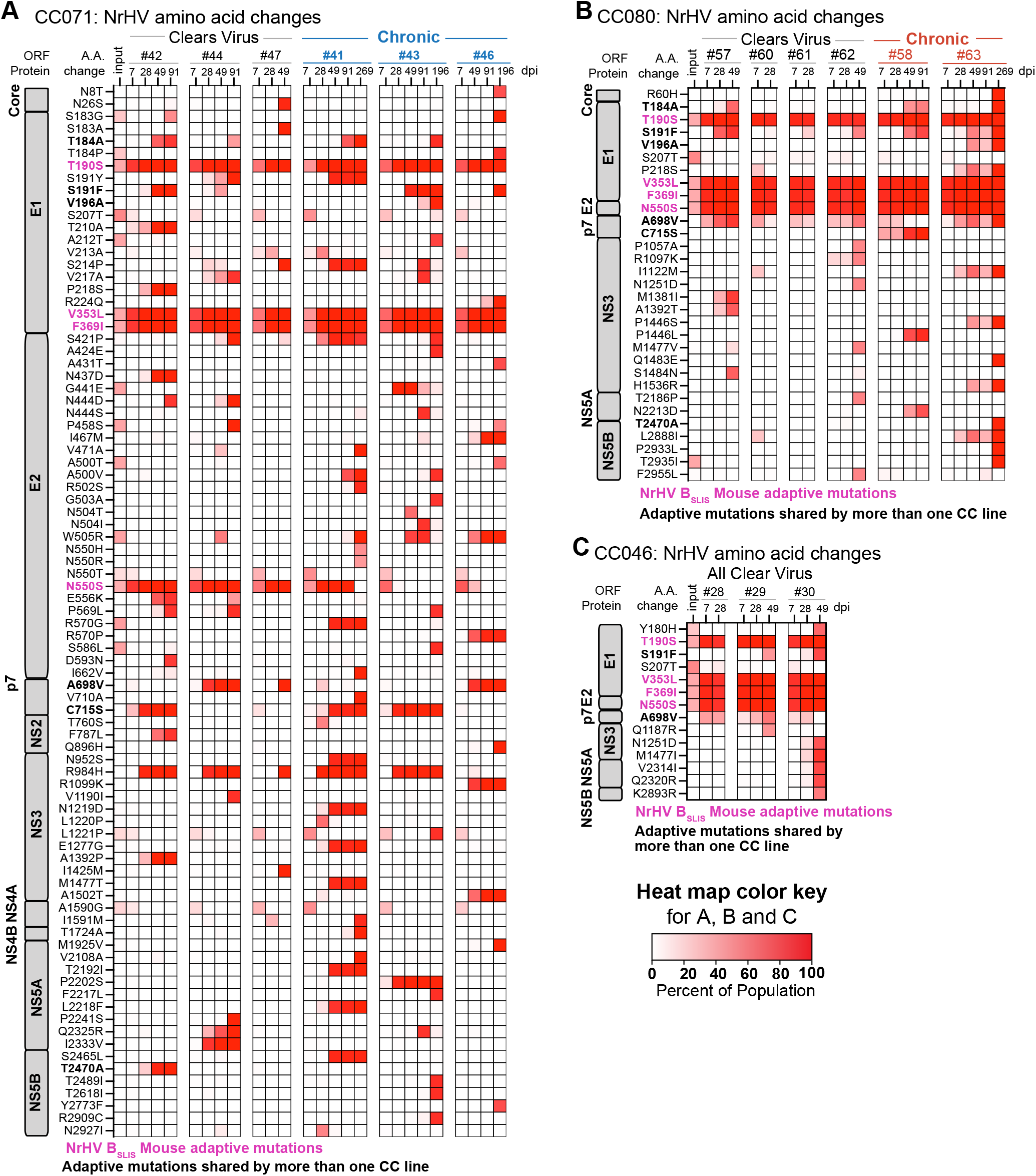
Host genetics drives differential viral evolution. Amino acid changes observed in the NrHV ORF for **(A)** CC071 animals that clear (#42, 44, 47) and become chronically infected (#41, 43, 46), **(B)** CC080 animals that clear (#57, 60, 61, 62) and become chronically infected (#58 and 63) and **(C)** CC046 animals that clear (#28, 29, 30). The heat map color intensity indicates mutation frequencies (%). Mouse adaptive mutations present in the inoculum (T190S, V353L, F369I, and N550S) are highlighted in hot pink. Changes shared by more than one CC strain are in bold.

### The pathologic hallmarks of chronic liver infection vary with time and host genetics

To determine the pathological consequences of acute and chronic NrHV infection, we evaluated hematoxylin and eosin-stained liver tissue sections. The hepatic architecture did not vary among mock infected WT, CC046, CC071 and CC080 mice (Fig. 7). For WT mice, the kinetics of liver pathology largely mirrored that of virus replication. On 2dpi, randomly distributed foci of inflammatory cells associated with single cell necrosis of hepatocytes were noted but by 7dpi, inflammation was notably increased with moderate lymphocytic inflammation predominantly focused on portal triads along with focal, random small clusters of inflammatory cells in the parenchyma all of which was largely resolved by 28dpi (Fig. 7 and S11). In infected CC046, a mild lymphocytic periportal inflammation was noted 7dpi, which became moderate by 28dpi and persisted although more mildly to the end of the study, 8 months after mice had cleared virus (Fig. 7 and S12). Although liver tissue sections from CC071 mice were unremarkable on 2dpi, by 7dpi, moderate periportal inflammation was observed in infected CC071 mice with increased lymphocytes in circulation (Fig. 7 and S13). By 28dpi, periportal and intraparenchymal inflammation was noted and hepatic nuclear inclusions (Fig. S13 Inset), known to be associated with liver disease such as hepatocellular carcinoma and non-alcoholic fatty liver disease, were observed ^22,23^. In addition, karyomegaly (i.e. enlargement of nuclei), known to be associated with abnormal liver function, was observed in CC071 infected mice at 28dpi (Fig. S13)^24^. Unlike WT and CC046, by the end of study, moderate periportal inflammation remained in infected CC071 mice. Like CC071, at 2dpi, liver sections from CC080 infected mice were unremarkable (Fig. 7 and S14). Over time, periportal inflammation and foci of intraparenchymal inflammation increased in frequency and severity which by the end of the study was like that in CC071. Lastly, we performed Masson’s Trichrome stain on liver tissue sections from the end of the study (i.e. 272 days post infection) to determine if the chronic infection and inflammation was associated with liver fibrosis (Fig. 8). Remarkably, early bridging fibrosis was observed in chronically infected CC071 mice but not in chronically infected CC080 mice suggesting that genetic factors guide multiple disparate pathologic outcomes.

**Figure 7.**
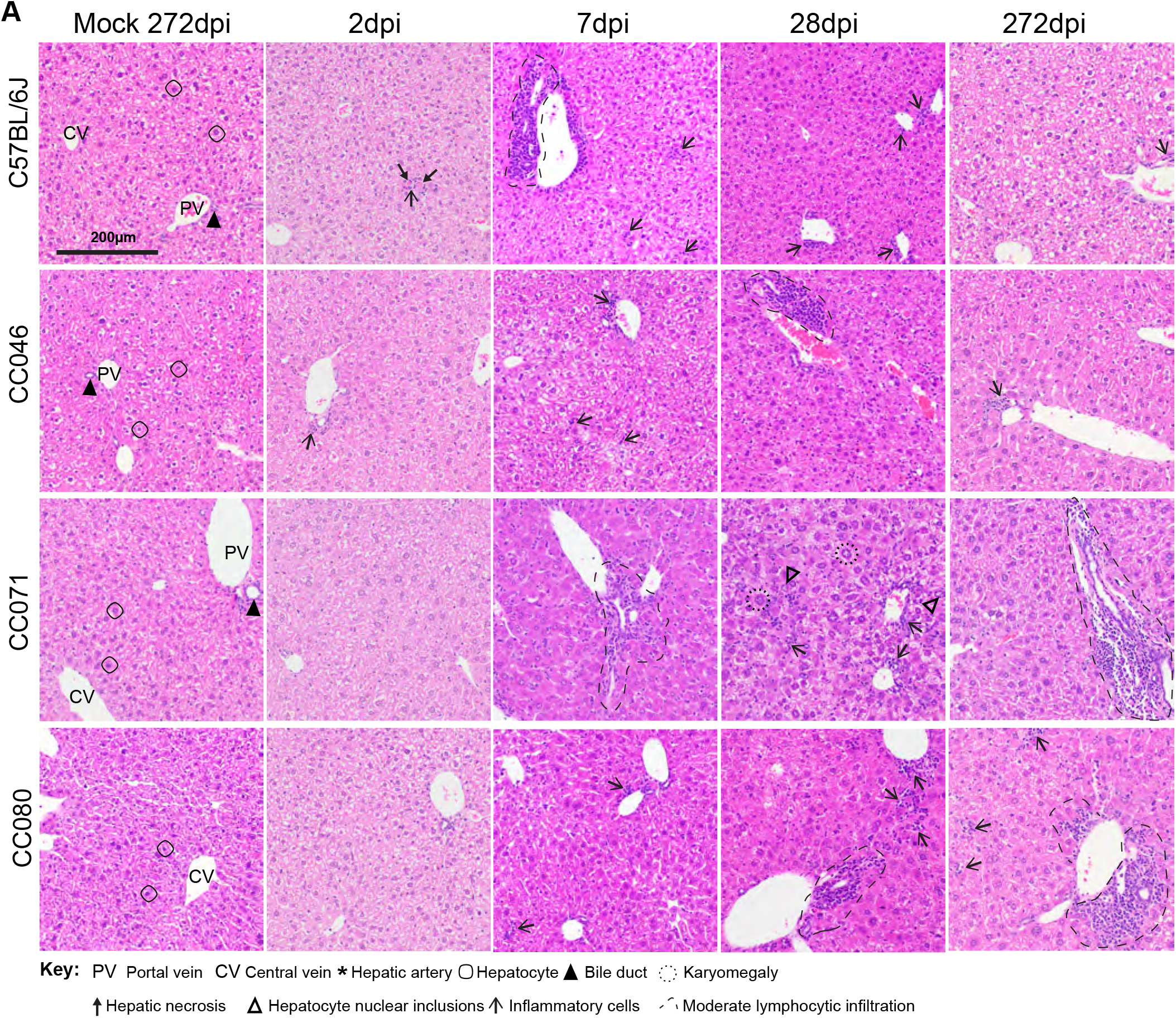
The pathologic hallmarks of chronic liver infection vary with time and host genetics. Liver tissue sections from mock or NrHV infected WT, CC046, CC071 and CC080 mice at 2, 7, 28 and 272 dpi were stained with hematoxylin and eosin. Hallmark features of normal liver architecture including portal vein, bile duct, hepatic artery, central vein and hepatocytes are noted in each mock panel.

**Figure 8.**
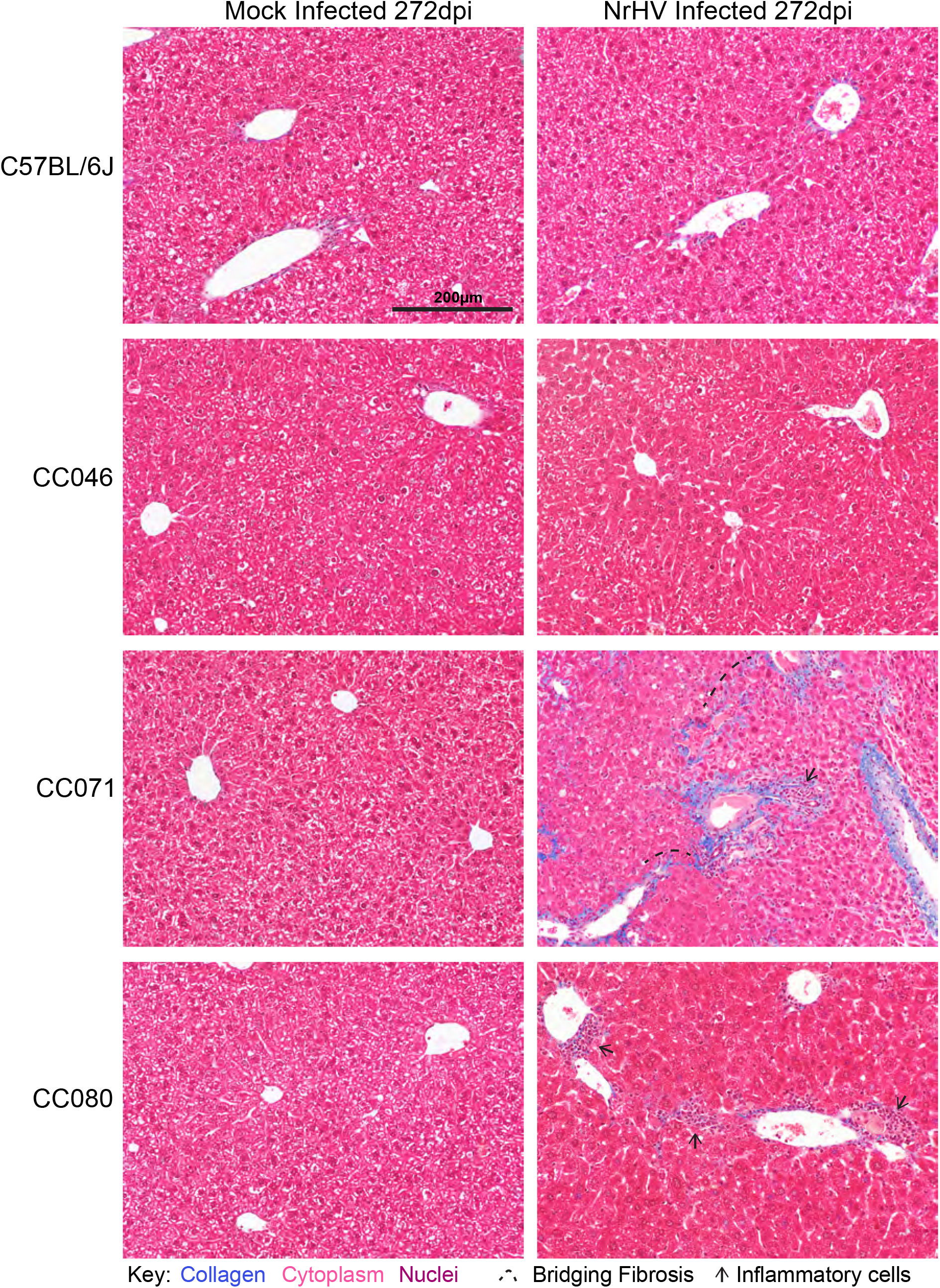
Liver fibrosis is associated with chronic liver infection. Liver tissue sections from mock or NrHV infected WT, CC046, CC071 and CC080 mice at 272 dpi were stained with Masson’s Trichrome which stains collagen blue. Inflammatory cells and early bridging fibrosis are noted.

## Discussion

Here we describe new mouse models of chronic hepacivirus infection that recapitulate multiple aspects of human HCV infection including high titer replication in the liver, persistent infection with virus evolution over time, chronic liver inflammation and liver fibrosis (Fig. S15)^25,26^. For hepacivirus research, the CC provides new models for the mechanistic study of acute and chronic infection, pathogenesis, immunity and vaccination and should provide insights into how host genetics affects acute and chronic infection at an unprecedented resolution.

Hepacivirus and hepatocyte interactions guide the trajectory of infection likely involving but not limited to the interplay of viral innate immune antagonists, antiviral innate immune sensors, innate (e.g. Kupffer cells) and adaptive immune cells (e.g. T-cells), cytokines and chemokines, hepatocyte turnover, and host genetic variation (e.g. *IL28B* SNPs)^27-30^. Here, we describe three genetically distinct mouse strains with a multitude of different hepacivirus infection phenotypes and disease trajectories. Interestingly, CC071 mice also have increased susceptibility to classical flaviviruses such as Zika virus (ZIKV), Powassan virus and West Nile virus but not to the bunyavirus Rift Valley fever virus indicating that this strain is particularly sensitive to the *Flaviviridae*^31-33^. JAK/STAT signaling, a key component of the interferon response, appears to function normally in CC071 as interferon treatment of MEFs from CC071 or WT results in similar levels of STAT1 phosphorylation suggesting that defects in JAK/STAT signaling is not driving increased flavivirus susceptibility^32^. Neither CC046 nor CC071 appear to have a broad immune deficiency as they are not more susceptible to *Salmonella* Typhimurium infection than WT mice (i.e. 129, 129S2/ SvPasCrl)^34^.

There is a dearth of small animal models of chronic hepacivirus infection within which to study the virus and host factors that contribute to pathogenesis, mechanisms of viral evolution and evasion of adaptive immunity. Sophisticated transgenic mice, human liver chimeric mice and even mice reconstituted with both human liver and immune cells have been employed to study HCV infection, but species incompatibility complicates the modeling of immunity and chronic disease^35^. Here, we provide an orthogonal approach with NrHV^9,10,36^. Although CD4 T-cell depletion prior to NrHV infection of WT mice can facilitate chronic infection^9^, using the CC, we provide a complementary approach demonstrating chronic infection and fibrosis in mice is possible without immune manipulation and that host genetic variation affects viral evolution. Viral evolution was greatest in CC071 mice, with the majority of change occurring in E1 and E2, presumably the main targets of the neutralizing antibody response. The MHC and T-cell related loci of WT, CC046, CC071 and CC080 are different suggesting the potential for differential adaptive immune responses. It is likely that acute resolving and chronic infection are complex processes involving infected cell intrinsic factors, and innate and adaptive immunity. Genetic differences in all these elements could in principle determine the differential outcomes observed between strains. Unfortunately, MHC haplotypes in CC strains remain largely unknown unlike common WT strains of mice.

In summary, we provide new small animal models of chronic hepacivirus infection, evidence that host genetic variation can profoundly affect the outcome of infection and a model genetic system within which to map the loci guiding virus clearance and chronicity. These models have the power to provide both basic insights into hepacivirus biology, hepatocyte biology, innate and adaptive immunity, mammalian genetics as well as new immunocompetent systems within which to study hepacivirus therapeutic efficacy and vaccine performance. Thus, these and future studies should be of broad interest to the field of infectious disease biology, hepatology and immunology.

## Materials and Methods

### Recombinant Virus Production

DNA from consensus clones NrHV-A, NrHV-B and the mouse adapted NrHV-B_SLIS_, carrying the T190S, V353L, F369I and N550S mutations (clones to be published elsewhere), were linearized using *MIuI* and purified using DNAclean and concentrator (Zymo). RNA was transcribed from 1 ug of *MIuI*-linearized DNA using T7 RiboMAX Express Large Scale RNA Production System (Promega). Template DNA was degraded using RQ1 DNAse for 30 min at 4°C and RNA was purified using RNAeasy mini kit (Qiagen). 4-week-old NRG mice were injected intra-hepatically using 10 µg of RNA in a maximum volume of 50 µL of PBS+RNA and one week post injection a terminal bleed was performed, and serum was pooled, titrated by qRT-PCR and stored in -80°C freezer. These passage zero virus stocks were amplified through intravenous infection of new 4-week-old NRG mice (N = 10/virus strain to be amplified) with 10^4^ genomic equivalents (GE) after which serum harboring the new passage #1 stocks were harvested, titrated and stored as above. All experiments were performed using passage #1 of each strain mixed in equal amounts (1:1:1).

### Mouse Infection Studies

All Collaborative Cross (CC) mice were obtained in 2015-2021 from the Systems Genetics Core Facility at the University of North Carolina at Chapel Hill^37^. For the initial CC-strain screen, 9–12-week-old female CC004/TauUnc, CC017/Unc, CC021/Unc, CC024/GeniUnc, CC040/TauUnc, CC044/Unc, CC045/GeniUnc, CC046/Unc, CC071/TauUnc and CC080/TauUnc were obtained N = 8/CC strain. Age and sex matched WT C57BL/6J mice were purchased from Jackson Labs. Mice were anesthetized with a mixture of ketamine/xylazine and then infected via retroorbital injection of 100µl containing 1×10^5^ genome equivalents (G.E.) of recombinant NrHV diluted in DPBS (Gibco) (N = 5/strain). As a negative control, 3 mice/strain were mock infected with DPBS. Mice were weighed daily for the first week of infection and then weekly thereafter. To measure virus replication in live mice, whole blood was collected via submandibular bleed, was allowed to clot at room temperature in snap cap tubes, centrifuged at 9,600 x g for 10 minutes and then serum was transferred to a new tube and stored at -80°C until analysis. Complete blood count was also performed using a Vetscan HM5c automated veterinary blood analyzer using whole blood collected into EDTA tubes. At the end of week 4, animals were euthanized by isoflurane overdose, whole blood and serum were collected and treated as above. Liver tissue was collected from each animal and stored in RNAlater (Thermo) at -80°C or in 10% formalin prior to processing for histology.

For the subsequent two follow up studies, 7–14-week-old female CC046/Unc (CC046) (N = 17 NrHV infected, 13 mock infected), CC071/TauUnc (CC071) (N = 28 NrHV infected, 19 mock infected) and CC080/TauUnc (CC080) (N = 15 NrHV infected, 14 mock infected) were purchased from the UNC Systems Genetics Core in January through March 2019. Age and sex matched WT C57BL/6J mice were purchased from Jackson Labs (N = 23 NrHV infected, N = 17 mock infected). Mice were infected and followed for the first six weeks as described above. Bleeds and weights then performed every other week for 8 weeks followed then by monthly assessments. These studies were terminated after 37-38 weeks, and blood and tissue were collected as noted above.

### RNA isolation, qRT-PCR and Illumina Deep Sequencing

For reverse transcription – quantitative polymerase chain reaction (qRT-PCR) quantitation of viral RNA in serum, RNA was isolated from 5µl of mouse serum using the RNA Clean & Concentrator -5 (Zymo) or High Pure Viral Nucleic Acid kit (Roche) kits. qRT-PCR was performed using 5µl of eluted RNA, TaqMan Fast Virus 1-step Master Mix (Applied Biosystems) and primer/probes targeting NS3 primers/probe (IDT)^11^. The sequence of the NrHV NS3 primers and probes were: Primer 1: AAGCGCAGCACCAATTCC, Primer 2: TACATGGCTAAGCAATACGG, Probe: /56-FAM/CTCACGTAC/ZEN/ATGACGTACGGCATG/3IABkFQ/. NrHV standard curve RNA (MEGAscript, Thermo) was produced by PCR amplification of NS3 downstream of a T7 promoter. Reactions were performed in LightCycler 480 Multiwell Plates (Roche) in a LightCycler 480 (Roche) using the following program: 50°C for 30 minutes, 95°C for 5 minutes, followed by 40 cycles of 95°C for 15 seconds, 56°C for 30 seconds, and 60°C for 45 seconds, then 40°C for 10 seconds. For qRT-PCR to quantitate NrHV RNA, ISG expression and mRNA sequencing in liver tissue, liver tissue was homogenized in TRIzol reagent (Thermo), and RNA was isolated according to protocol. 500ng total RNA was utilized for each qRT-PCR reaction. ISG MX dynamin-like GTPase 1 (Mx1, Thermo Fisher, Mm00487796_m1) expression was assessed in select samples. Relative expression of Mx1 to housekeeping gene glyceraldehyde-3-phosphate dehydrogenase (GAPDH, Thermo Fisher, Mm99999915_g1) was calculated by ΔΔCT method^38^. Illumina total RNA-seq was performed by the UNC High Throughput Sequencing Facility (HTSF). Stranded mRNA libraries were prepared (Kapa mRNA) and read on the Illumina HiSeq4000 platform (single end 1×50 read length). For analysis of raw reads, sequenced reads were mapped to the mouse reference transcriptome (Ensemble; Mus musculus version 108) using Kallisto (version 0.46.0). Transcript quantification data was normalized using the TMM method in EdgeR (version 3.38.4) and differentially expressed genes (p.Adj.val < 0.05; logLC > 1.5) were identified using linear modeling with limma (version 3.52.2) using R (version 4.2.0) in R Studio (version 2022-04-19). Gene ontology analysis was carried out using the Ingenuity Pathway Analysis (Qiagen). Heat maps were generated using Broad Institute’s Morpheus web application (https://software.broadinstitute.org/morpheus/). Hierarchical clustering was performed by One Minus Pearson Correlation with Average Linkage clustering by columns and rows. Sequence data is available on the Gene Expression Omnibus (GEO, accession # GSE###).

### Liver Pathology and Digital Quantitation of Viral RNA in Liver Tissue Sections

To assess liver pathology, we formalin fixed, paraffin embedded liver tissues, generated sections (∼5µm) and stained then with hematoxylin/eosin (Richard-Allen Scientific) on an Autostainer XL from Leica Biosystems. To assess liver fibrosis, sections were stained using Masson’s Trichrome (Richard-Allen Scientific). For CD4 immunohistochemical (IHC) staining, antigen retrieval (20 minutes at 100°C in Leica Bond-Epitope Retrieval Solution 2 pH-9) was performed on dewaxed tissues prior to labeling with CD4 antibody (ab183685, Abcam) at 1:2,000 for 1h followed with Novolink Polymer (RE7260-CE) secondary antibody labeling on a Leica Bond III Autostainer. Antibody labeling was detected using 3,3’-diaminobenzidine (DAB) and the Bond Intense R detection system (DS9263). Tissue sections were evaluated by a Board Certified Veterinary pathologist.

NrHV viral RNA was visualized in liver tissue sections by *in situ* hybridization (ISH) using RNAscope probes (Item # V-NrHV-PP) designed by Advanced Cell Diagnostics based on the whole genome sequence of NrHV (Genebank Accession # MF113386.1). ISH on liver tissue sections was performed on a Leica Rx autostainer (Leica Biosystems) with a hematoxylin counterstain. For quantitation, slides were scanned on a Versa slide scanner (Leica Biosystems) with a 40X power objective and a Point Gray camera (8-bit image at 0.137152 microns/pixel) and imported to Definiens Architect XD 2.7 for analysis with Tissue Studio version 4.4.2. Per tissue section per animal, total tissue area was calculated and total numbers of cells per section was calculated using the nuclear counterstain as a guide. Similar numbers of cells were assessed in each tissue section per timepoint. NrHV positive cells were identified by ISH positivity. The spot size threshold was 1.5 µm². Spot areas were ranked as None/Low (2 µm²), Low/Medium (6 µm²), or Medium/High (10 µm²). Resultant data was used to determine the infection frequency per tissue section.

### Deep sequencing NrHV genomes

Serum from NrHV infected mice was diluted in PBS (25 µL in 225 µL PBS), added to a 2 mL Phasemaker tube (Thermo Fisher Scientific), mixed with 750 µL TRIzol LS Reagent (Thermo Fisher Scientific) after which 200 μl chloroform was added, shaken for 15 seconds, incubated for 3 min at room temperature, and centrifugated at 12,000g for 15 min at 4°C. The aqueous phase was removed and mixed with 450 μl ethanol and transferred to an RNA Clean & Concentrator-5 column (Zymo Research) for downstream RNA purification and concentration. Reverse-transcription (RT) was performed with Maxima H Minus Reverse Transcriptase (Thermo Fisher Scientific). Samples were pre-incubated in the presence of RNase inhibitors (Promega) at 65°C for 2 min prior to addition of the RT enzyme, followed by incubation at 50°C for 2h using 0.1 μM primer TS-O-00319 (GCTTCCTGGAGCGGGCTAGATACTG). Amplification of the complete ORF was performed using Q5 Hot Start High-Fidelity DNA Polymerase (New England Biolabs), including high GC Enhancer, and the primer pair Reverse TS-O-00318 (CCAAGCCCCAATGCCGTCCGGCACCGCTGCCCTTTTCGG) and Forward TS-O-00361 (AGGTGAAGGGGGCATCGATG) (0.5 μM each). The PCR cycling parameters were 98°C for 30 s, followed by 37 cycles of 98°C for 10 s, 65°C for 10 s and 72°C for 8 min, with a final extension at 72°C for 10 min. Amplified DNA was purified using DNA Clean & Concentrator-25 columns (Zymo Research), and libraries for deep sequencing were prepared using the NEBNext Ultra II FS DNA Library Prep Kit for Illumina (New England Biolabs). Quality of DNA libraries was validated using a 2100 Bioanalyzer Instrument (Agilent). DNA libraries were quantitated via Qubit dsDNA High-Sensitivity Assay Kit (Thermo Fisher Scientific) and libraries were pooled in equimolar concentrations prior to denaturation. Pooled libraries were loaded on a MiSeq v3 150 cycle flow-cell, and sequencing performed on a MiSeq benchtop sequencer (Illumina). Analysis was performed as previously described^10,39^, In short, pair-end reads were trimmed and filtered by Sickle and subsequently mapped by BWA onto the NrHV-B reference (GenBank: ON758386) using the MEM algorithm. Samtools were used to process the alignment files and Lofreq were applied to call SNPs that were translated by SNPEffect. Multiple mutations in the same specific codons were subsequently resolved by LinkGE. Pairwise distance and dN/dS ratios were calculated form the vcf files by SNPGenie.

## Author Contributions

TPS, MTF, CMR, JB and TKHS designed the studies. AJB, JW, RW, UF, NC, AW, FRM, MTF, CLL, SRL, CN, GDLC, BRM, YX, SAM, and TPS performed the studies. EB, MNB and CMR provided recombinant NrHV for these studies. TPS, RW, UF, EB, SAM, TKHS, JB and CMR wrote and edited the manuscript. All authors read and approved this manuscript.

## Acknowledgements

This work was funded by R01 grant from the National Institute of Allergy and Infectious Disease (NIAID, AI131688 to C.M.R.), a U19 grant from NIAID (U19AI100625 to Ralph Baric and Mark Heise), the Independent Research Fund Denmark (1030-00426 to Troels K. H. Scheel), Advanced Grant 4004-00598 to Jens Bukh), the Danish Cancer Society (R204-A12639 to Jens Bukh and Troels K. H. Scheel), the Novo Nordisk Foundation (Distinguished Investigator Grant NNF19OC0054518 and Tandem NNF19OC0055462 to Jens Bukh), and the European Research Council (Starting Grant 802899 to Troels K. H. Scheel). Raphael Wolfisberg. was supported by an Early Postdoc Mobility Fellowship (P2BEP3_178527) and a Post-doc Mobility Fellowship (P400PB-183952) from the Swiss National Science Foundation. We gratefully acknowledge the technical support from the UNC High Throughput Sequencing Facility. This facility is supported by the University Cancer Research Fund, Comprehensive Cancer Center Core Support grant (P30-CA016086), and UNC Center for Mental Health and Susceptibility grant (P30-ES010126). We would like to thank Louise B. Christensen (Department of Clinical Microbiology, University of Copenhagen) for laboratory assistance and the Department of Clinical Microbiology, Hvidovre Hospital for access to Illumina miSeq equipment. Collaborative Cross (CC)^37^ mice were obtained in 2017-2021 from the Systems Genetics Core Facility at the University of North Carolina at Chapel Hill. We would also like to thank Dr. Amit Kapoor from the University of Ohio and Dr. Brad Rosenberg from Mt. Sinai Medical School for helpful and insightful discussion of this work.

## Supplemental Figures

**Figure S1.**
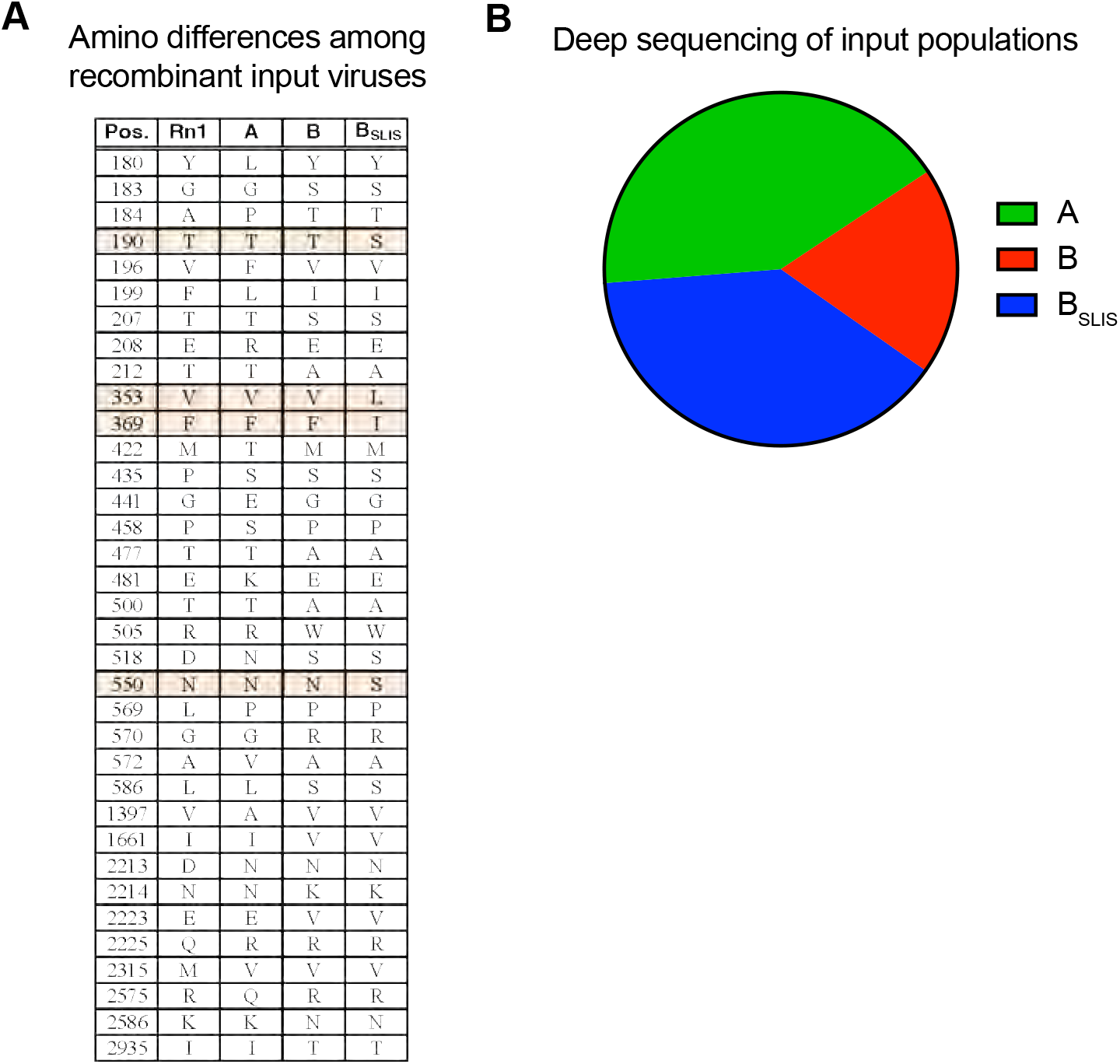
Deep sequencing of recombinant virus populations utilized for *in vivo* studies. Associated with main Figure 1. **(A)** The amino acid sequence differences for NrHV-A (Genbank: MF113386), NrHV-B (Genbank ON758386) and B_SLIS_ (NrHV-B with the additional changes T190S, V353L, F369I and N550S) as compared to the RHV-rn1 reference isolate (Genbank: KX905133) virus strain. **(B)** Deep sequencing of the input virus pool which revealed a stoichiometry of 42% A, 19% B, 39% B_SLIS_. Three additional mutations (207T, 1221P and 1590G) were noted in all B virus sequences likely acquired during stock generation *in vivo*.

**Figure S2.**
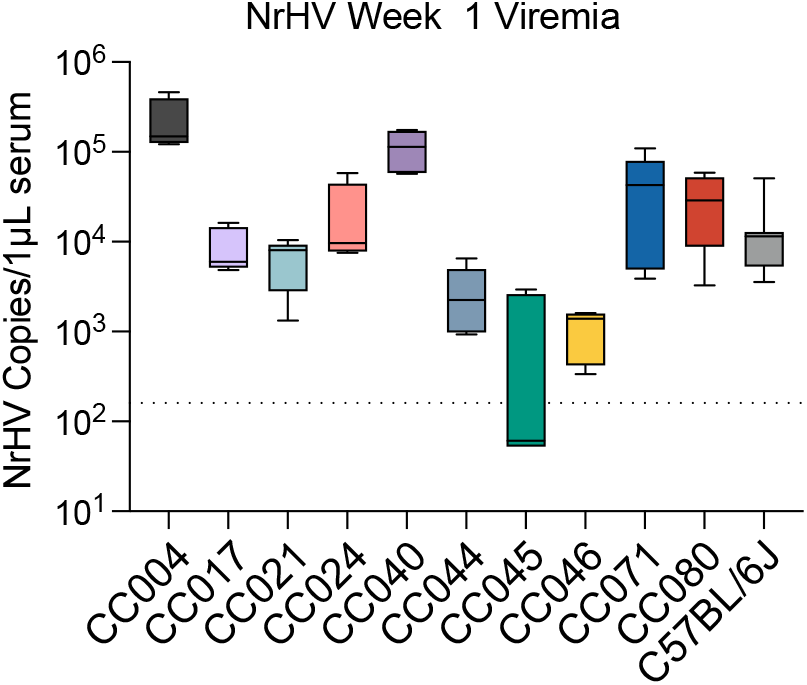
NrHV week 1 viremia for 10-strain Collaborative Cross screen. Associated with main Figure 1. Ten CC strains and control C57BL/6J female mice 9-13 weeks in age were infected with 1×10^5^ G.E. of recombinant NrHV or negative control PBS via retroorbital injection and bled weekly to monitor viremia. Infected mouse numbers per strain were: C57BL/6J N = 8, CC004 N = 4, CC017 N = 4, CC021 N = 4, CC024 N = 5, CC040 N = 5, CC044 N = 5, CC045 N = 5, CC046 N = 6. CC071 N = 5, CC080 N = 4. For all strains PBS mock infected N = 3. Levels of viral RNA in 1µl serum as measured by qRT-PCR is shown.

**Figure S3.**
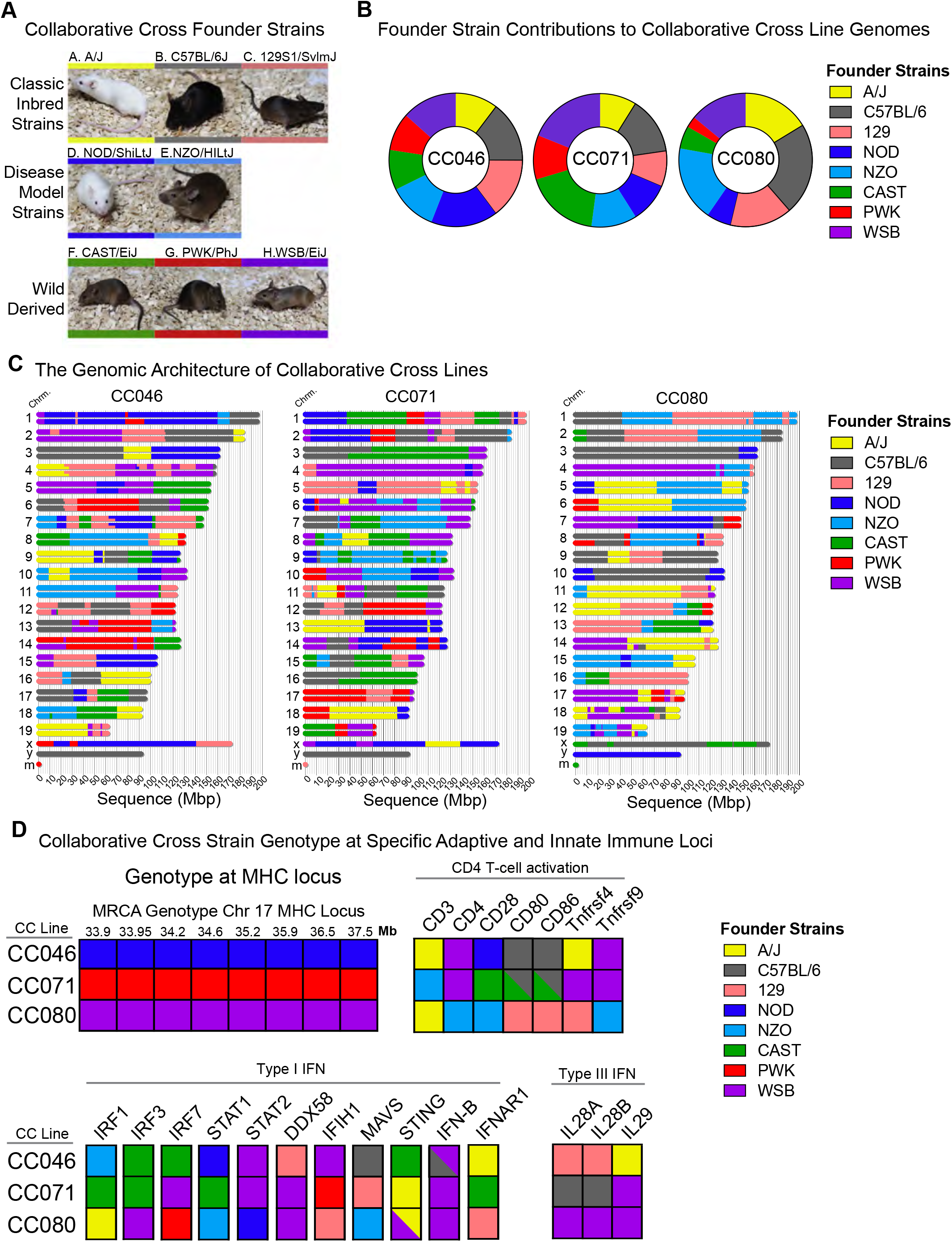
The genetic architecture of Collaborative Cross strains CC046, CC071, and CC080. Associated with main Figure 2. Figure adapted from Leist et al.^40^ **(A)** The CC is a recombinant inbred mouse reference population generated by breeding five classic inbred strains (A/J, C57BL/6, 129, NOD and NZO) with three wild strains (CAST, PWK and WSB). **(B)** The proportion of each founder strain comprising each CC strain of interest. **(C)** The genomic architecture of CC046, CC071 and CC080. The color segments of each chromosome pair correspond to loci donated by founder strains. **(D)** The genotype of CC046, CC071 and CC080 at the MHC locus, genes related to T-cell activation and type I and III interferon. The colors in the grid correspond to the genotype at each region.

**Figure S4.**
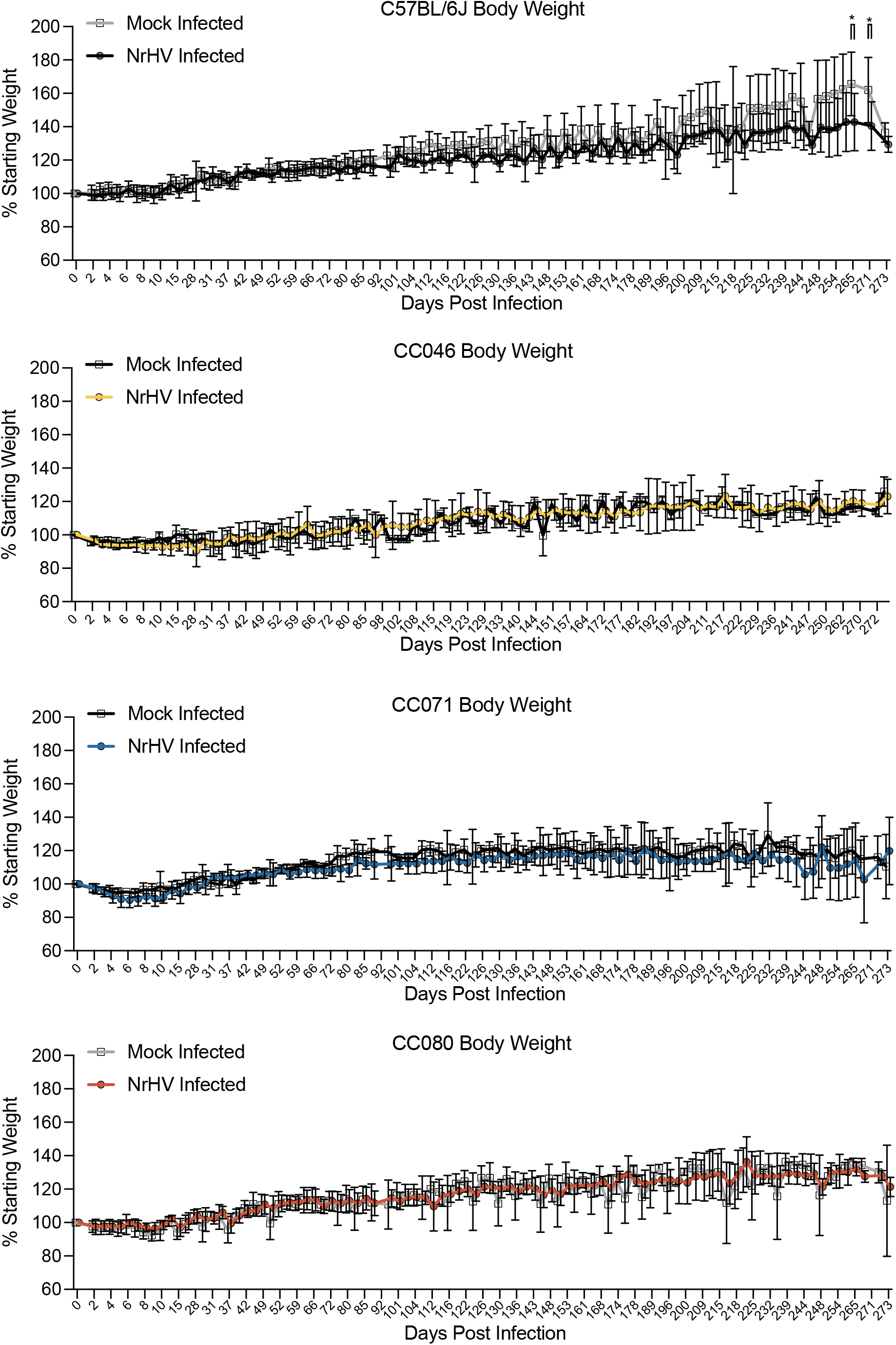
Body weight for mock and NrHV infected mice. Associated with main Figure 2. C57BL/6J, CC046, CC071 and CC080 mice were infected with 1×10^5^ G.E. of recombinant NrHV or negative control PBS via retroorbital injection. Body weights were measured daily for the first two weeks and then intermittently for the duration of the study to 272 or 273dpi. The percent starting weight is shown for age and strain matched mock PBS infected and NrHV infected over time.

**Figure S5.**
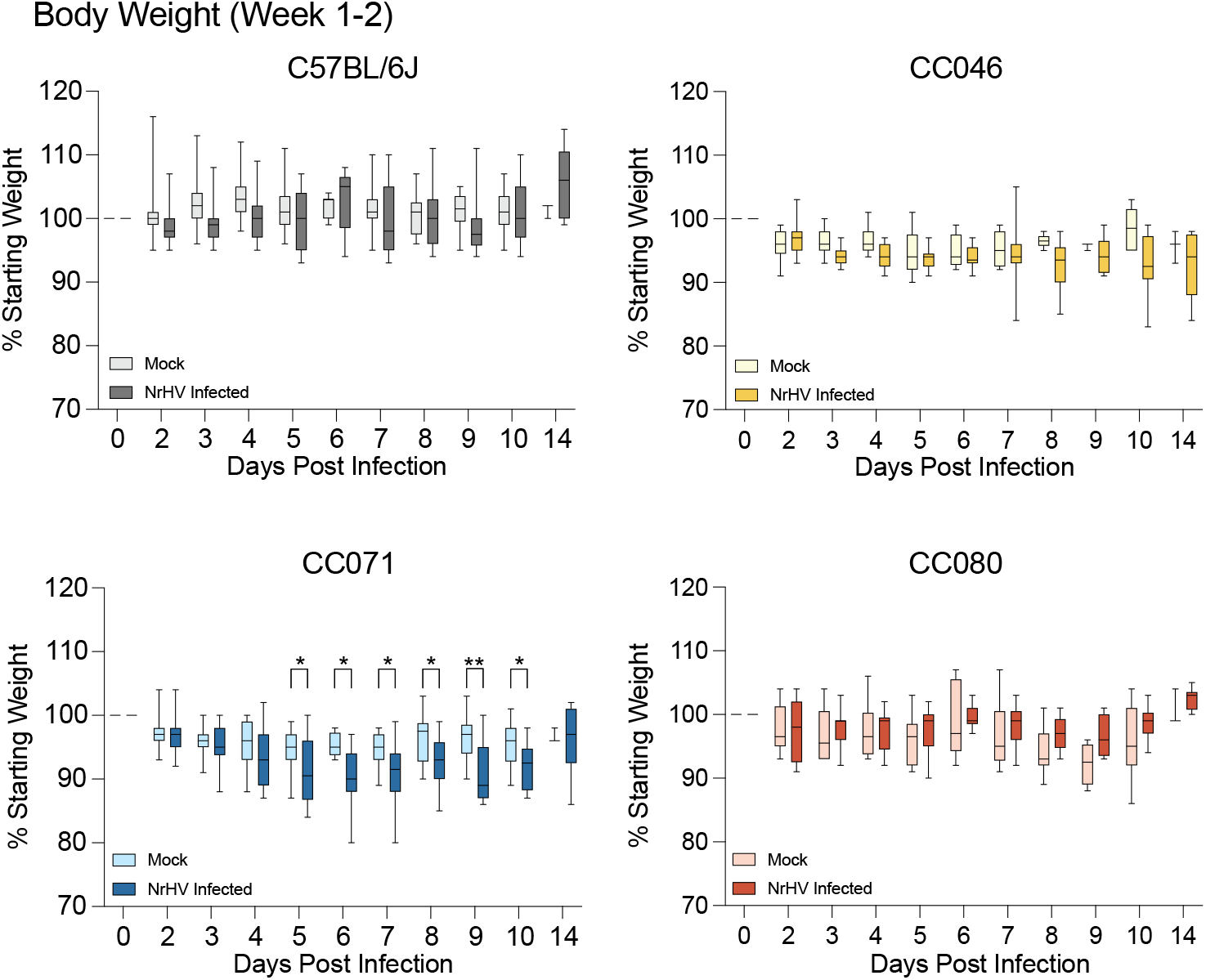
Body weight for mock and NrHV infected mice for days 0-14. Associated with main Figure 2 and Figure S4. C57BL/6J, CC046, CC071 and CC080 mice were infected with 1×10^5^ G.E. of recombinant NrHV or negative control PBS via retroorbital injection. Daily body weights are shown for the first 14 days of infection. Asterisks indicate statistical significance by two-way ANOVA with a Sidak’s multiple comparison test.

**Figure S6.**
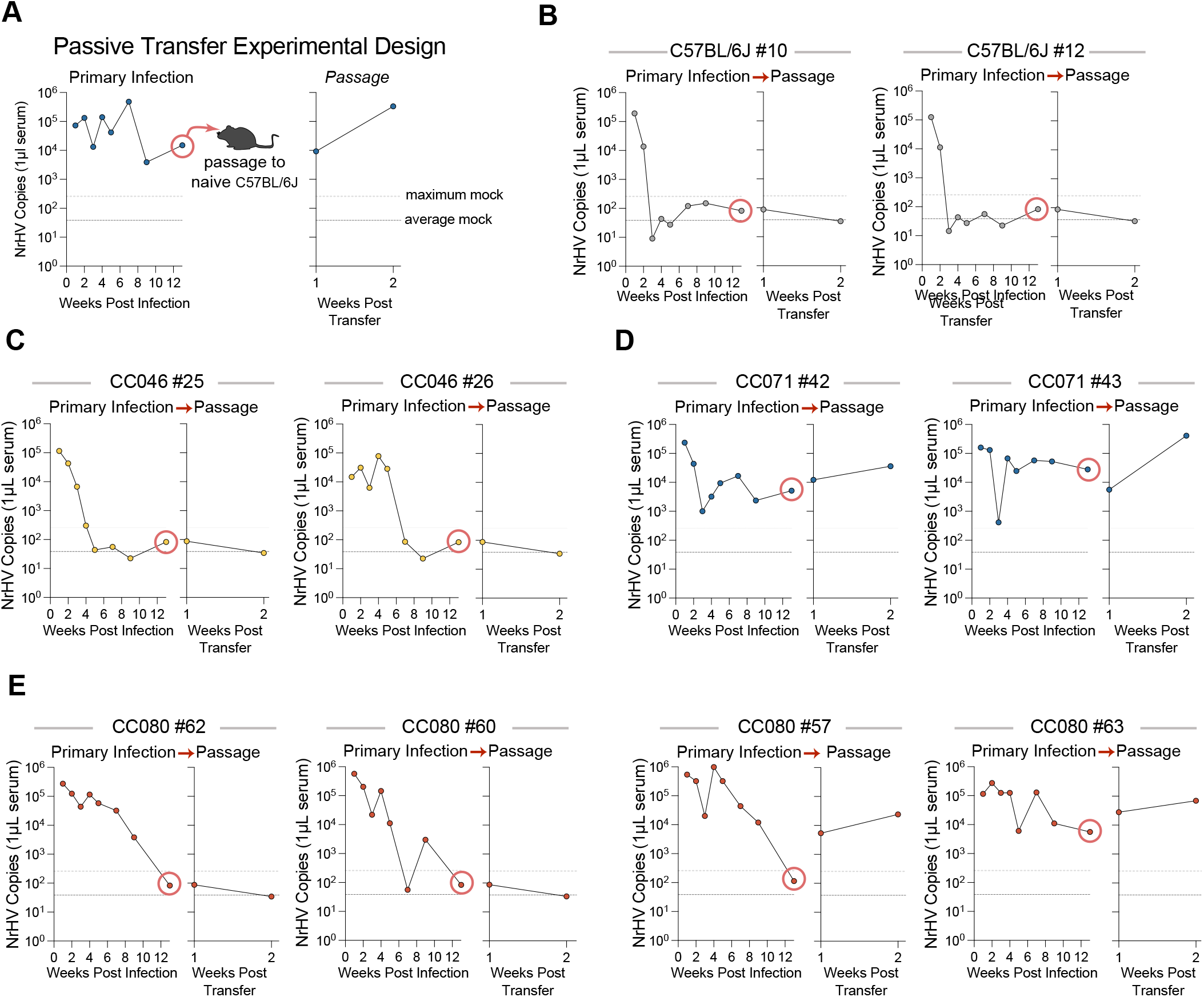
Passive transfer to demonstrate the presence of infectious virus. Associated with main Figure 2. **(A)** To demonstrate the presence of infectious virus in NrHV RNA positive sera, 5µl of serum from 12 weeks post infection was diluted in 150µl PBS and 100µl of this mixture was used to inoculate naïve C57BL/6J mice via the retroorbital route. **(B)** Transfer of serum from previously infected aviremic C57BL/6J mice to recipient naïve C57BL/6J mice. **(C)** Transfer of serum from previously infected but aviremic CC046 mice to recipient naïve C57BL/6J mice. **(D)** Transfer of serum from viremic CC071 mice to recipient naïve C57BL/6J mice. **(E)** Transfer of serum from three aviremic mice and one viremic CC080 mouse to three recipient naïve C57BL/6J mice.

**Figure S7.**
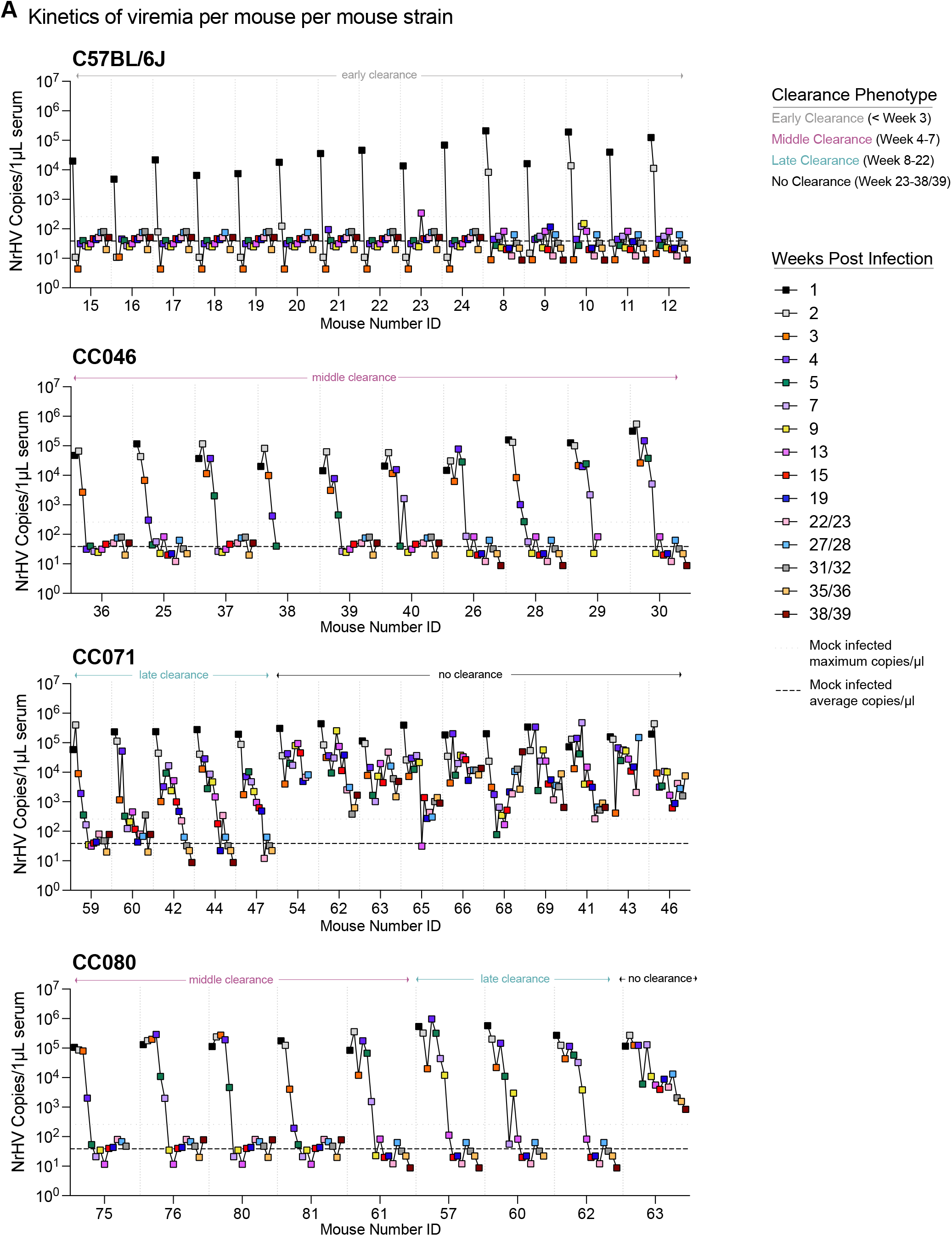
Kinetics of viremia in individual mice with different viral clearance phenotypes. Associated with main Figure 2. The levels of viral RNA in serum as measured by qRT-PCR are shown for C57BL/6J, CC046, CC071 and CC080. Four clearance phenotypes were noted: Early Clearance (< 3 weeks), Middle Clearance (Week 4-7), Late Clearance (Week 8-22), No Clearance (viremic at the end of study week 38/39).

**Figure S8.**
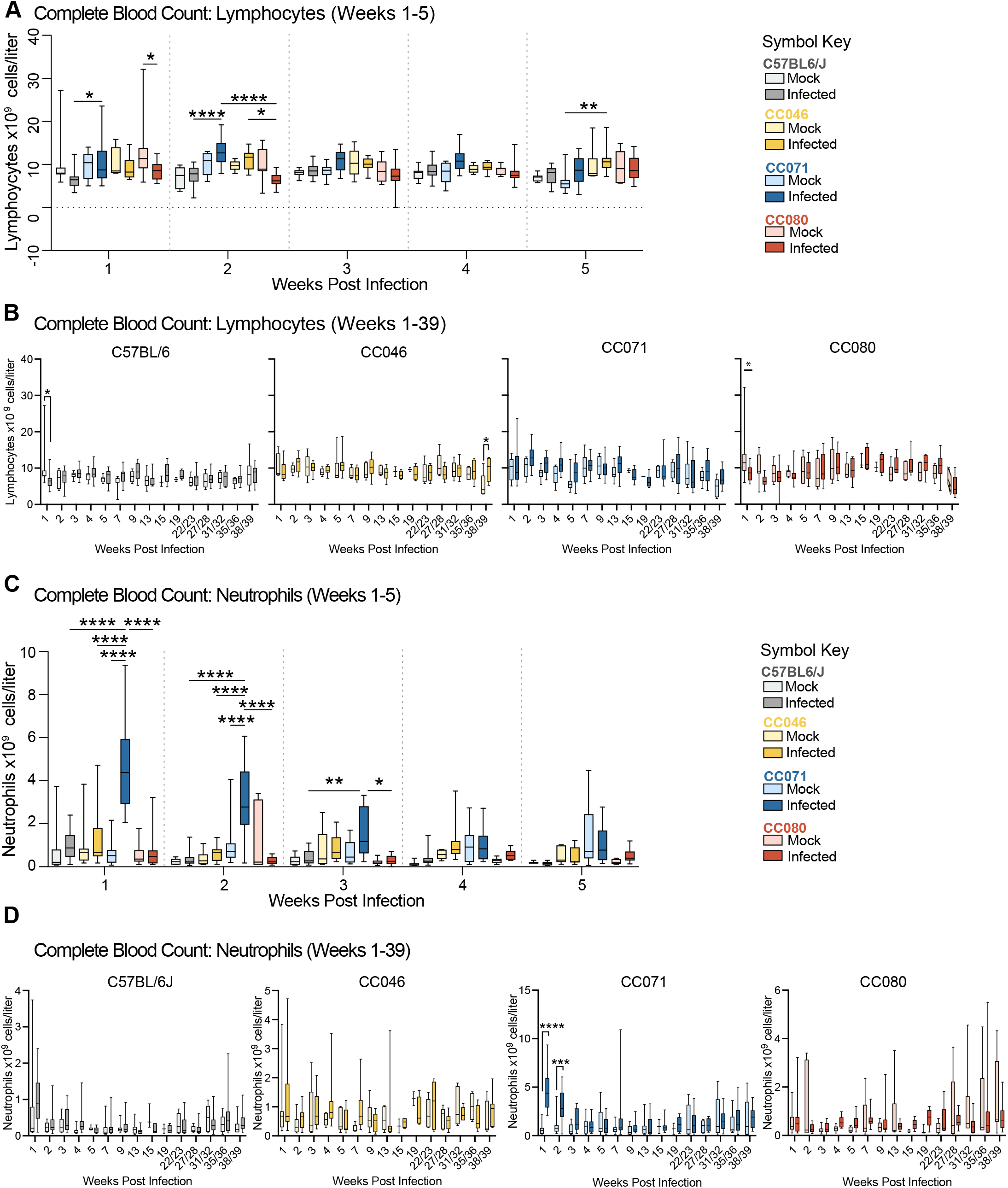
Total blood lymphocytes and neutrophils for all times post infection. Associated with main Figure 4. Complete blood count was performed using a Vetscan HM5 automated blood analyzer on whole blood isolated from mock and infected mice for the duration of studies described in Fig. 2. **(A)** Blood lymphocytes are shown for the first five weeks of infection or **(B)** for the duration of the study for each mouse strain. **(C)** Complete blood count for neutrophils for the first five weeks of infection or **(D)** for the duration of the study for each mouse strain. Asterisks indicate statistically significant differences by Two-Way ANOVA Tukey’s multiple comparisons test.

**Figure S9.**
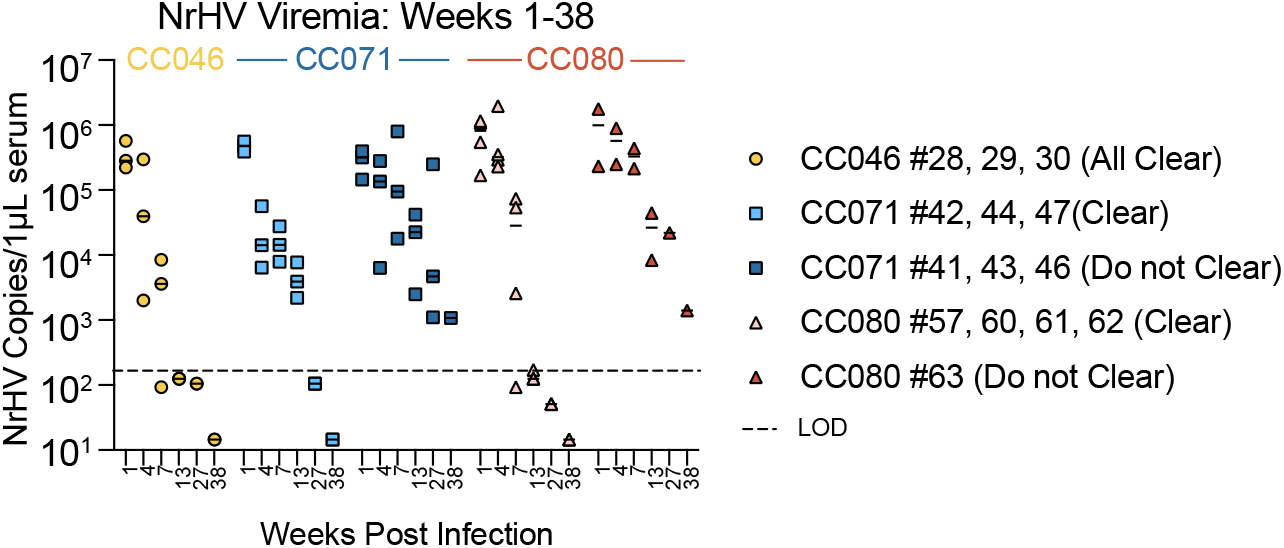
Viremia phenotypes for mice on which viral ORF sequencing was performed. Associated with main Figure 6 and Figure S12. The levels of viral RNA in serum as measured by qRT-PCR are shown for C57BL/6J, CC046, CC071 and CC080.

**Figure S10.**
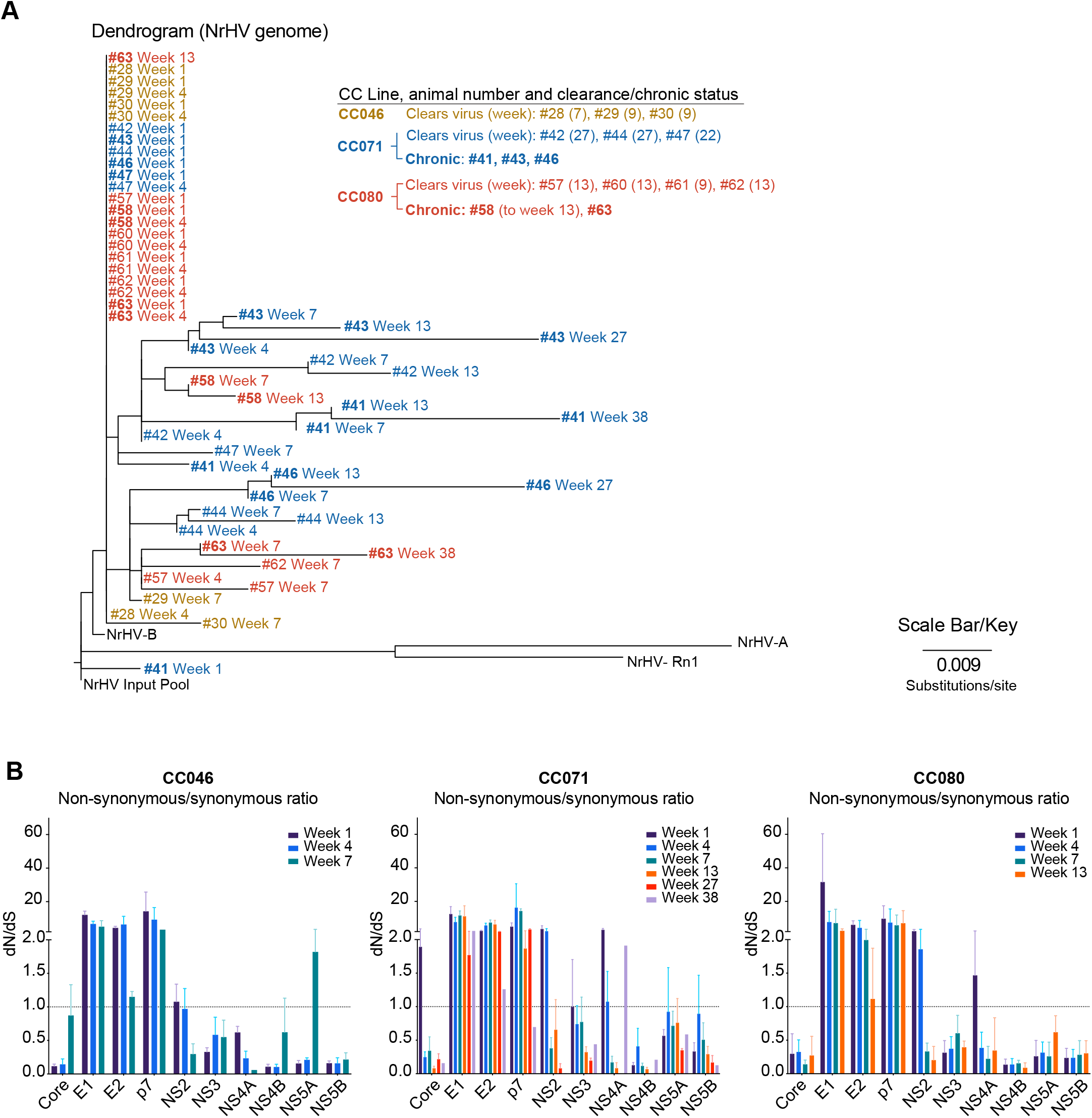
Chronic infection is associated with increased viral evolution. **(A)** Associated with main Figure 6 and Figure S9. The genetic relationships of virus populations isolated from NrHV infected CC046 (N = 3), CC071 (N = 6) or CC080 (N = 6) over time are shown in a neighbor joining tree created from whole viral ORF sequences from 1, 4, 7, 13, 27 and 38 weeks post infection. Infected animals and phenotypes are described in Figure 2. The viremia phenotypes per mouse are described in Figure S9. The dendrogram was generated by aligning full ORF sequences by MAFFT and building the phylogeny by maximum likelihood using PhyML using the general time reversible substitution model and rooted the inoculum input of the current study and the rat inoculum used for NRG mouse adaptation (NrHV-A; Genebank MF113386)^9^; NrHV-B (Genebank ON758386) and RHV-rn1 (Genebank KX905133)^36^ as outgroups. **(B)** Non-synonymous/synonymous (dN/dS) mutation ratios. Using whole ORF sequencing data from (A), dN/dS ratios were generated for each viral protein. Increased dN/dS is associated with increased evolutionary change.

**Figure S11.**
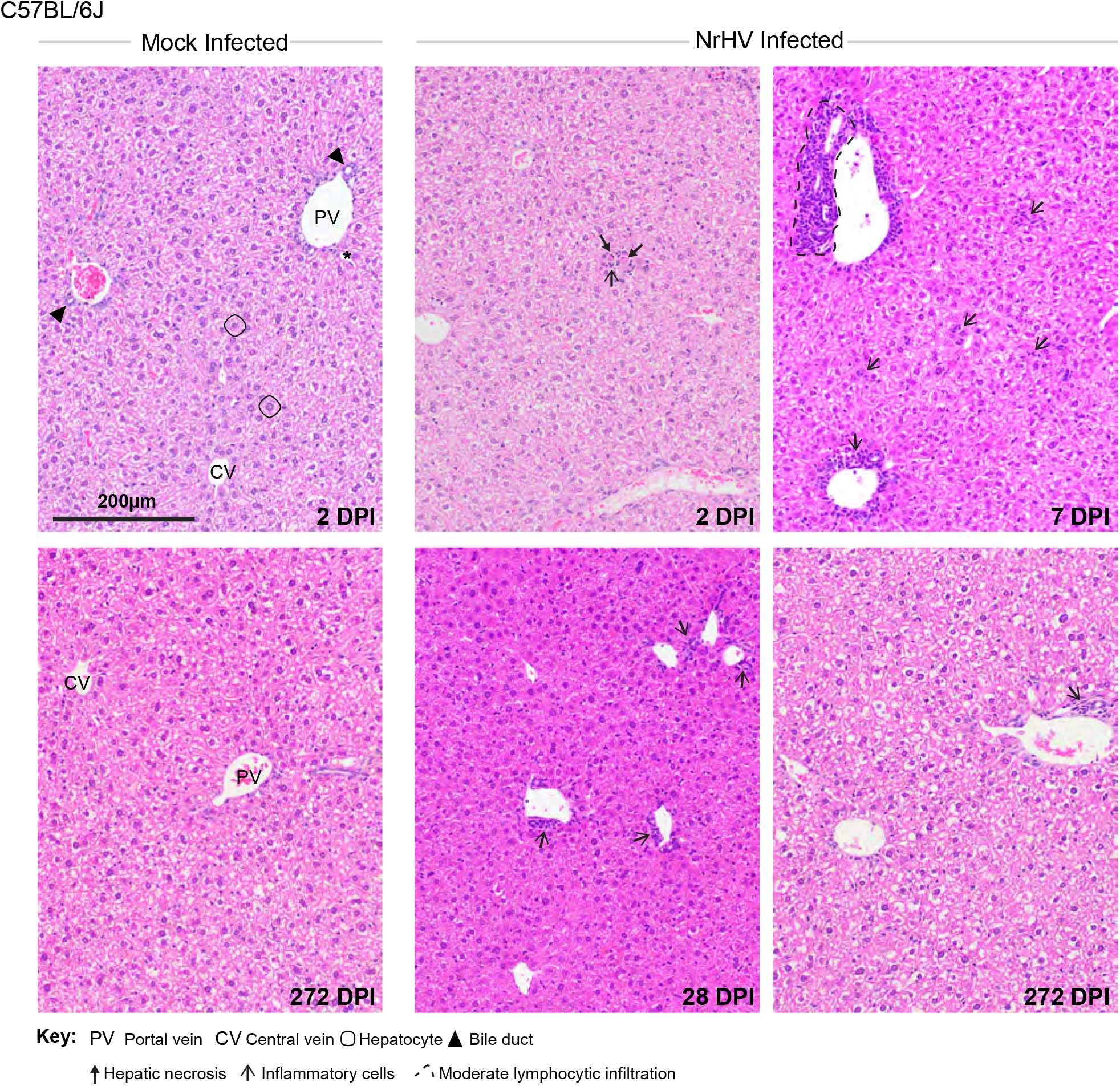
Liver pathology in mock and NrHV infected WT C57BL/6J mice. Liver tissue sections from mock or NrHV infected mice at 2, 7, 28 and 272 dpi were stained with hematoxylin and eosin. Hallmark features of normal liver architecture including portal vein, bile duct, hepatic artery, central vein and hepatocytes are noted in each mock panel. Other pathologic features are noted in the key.

**Figure S12.**
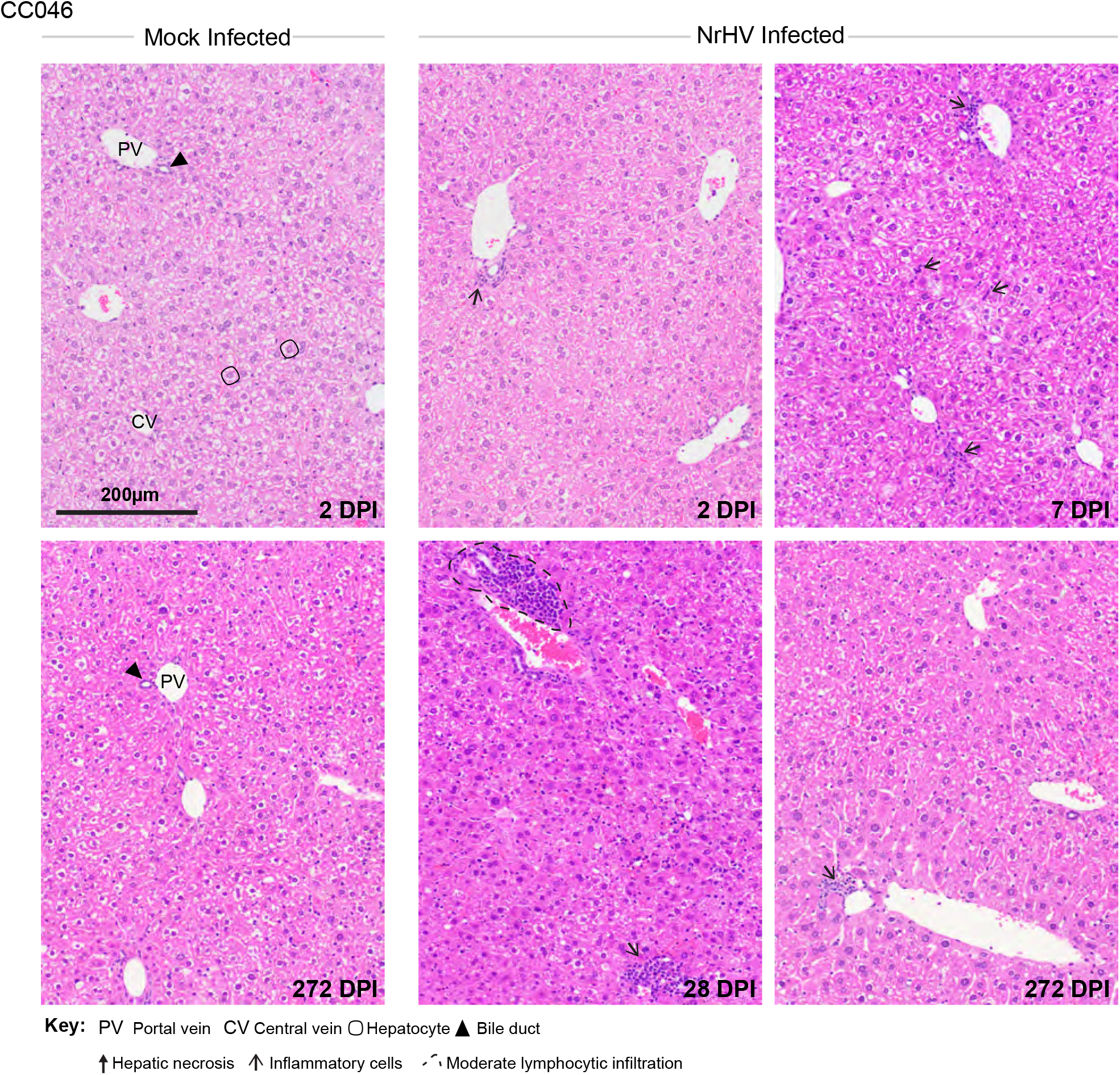
Liver pathology in mock and NrHV infected CC046 mice. Liver tissue sections from mock or NrHV infected mice at 2, 7, 28 and 272 dpi were stained with hematoxylin and eosin. Hallmark features of normal liver architecture including portal vein, bile duct, hepatic artery, central vein and hepatocytes are noted in each mock panel. Other pathologic features are noted in the key.

**Figure S13.**
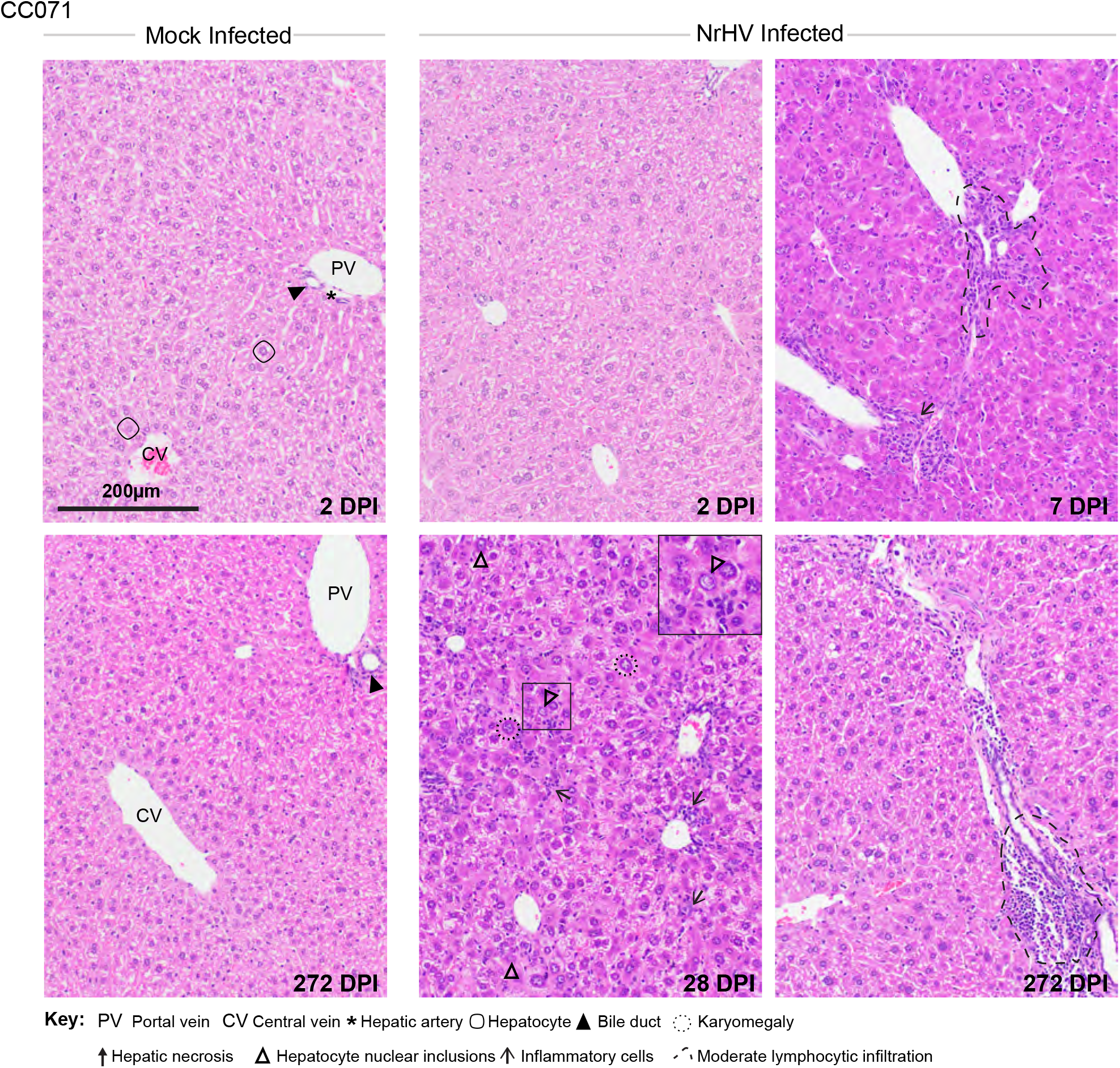
Liver pathology in mock and NrHV infected CC071 mice. Liver tissue sections from mock or NrHV infected mice at 2, 7, 28 and 272 dpi were stained with hematoxylin and eosin. Hallmark features of normal liver architecture including portal vein, bile duct, hepatic artery, central vein and hepatocytes are noted in each mock panel. Other pathologic features are noted in the key.

**Figure S14.**
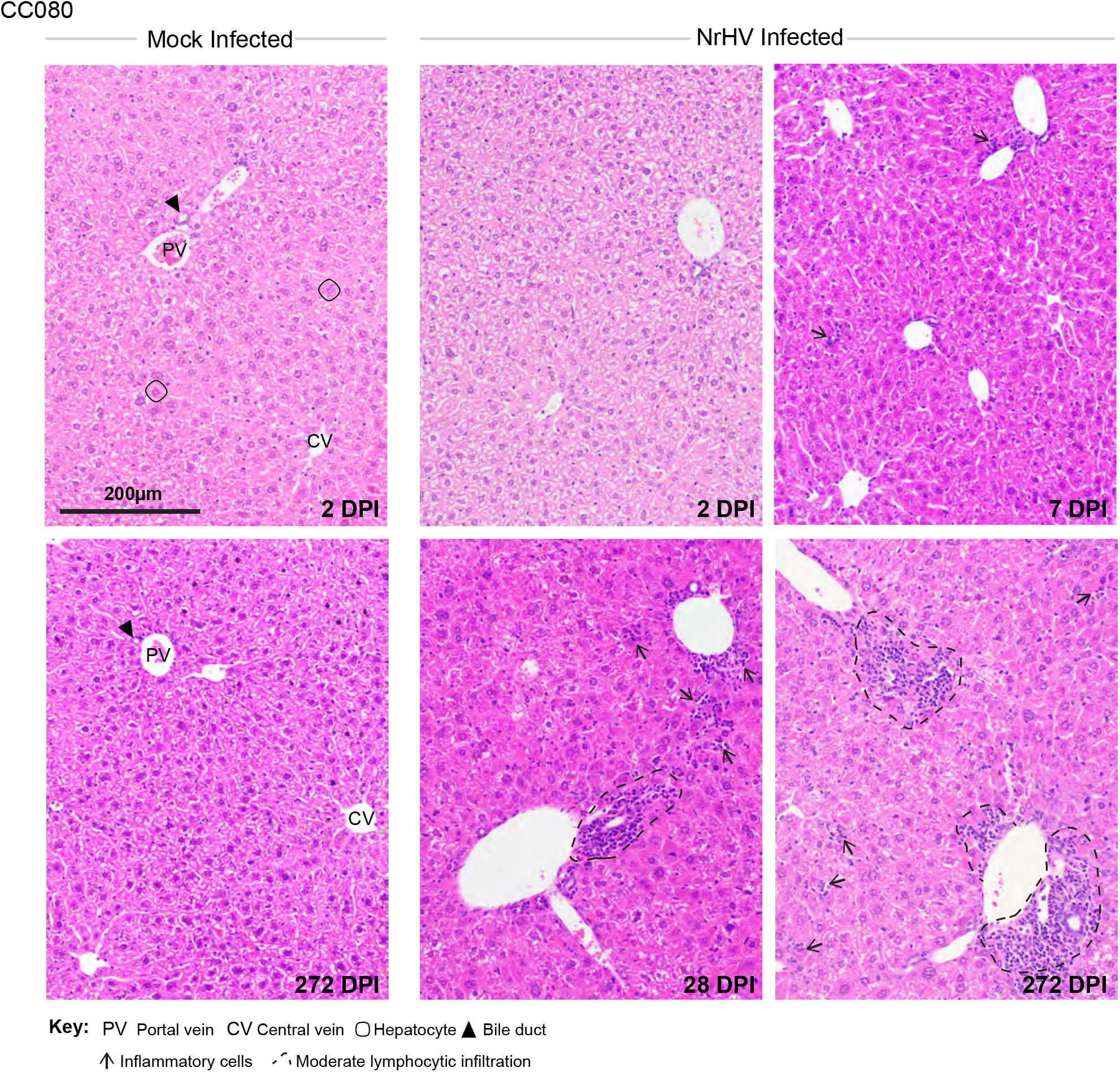
Liver pathology in mock and NrHV infected CC080 mice. Liver tissue sections from mock or NrHV infected mice at 2, 7, 28 and 272 dpi were stained with hematoxylin and eosin. Hallmark features of normal liver architecture including portal vein, bile duct, hepatic artery, central vein and hepatocytes are noted in each mock panel. Other pathologic features are noted in the key.

**Figure S15.**
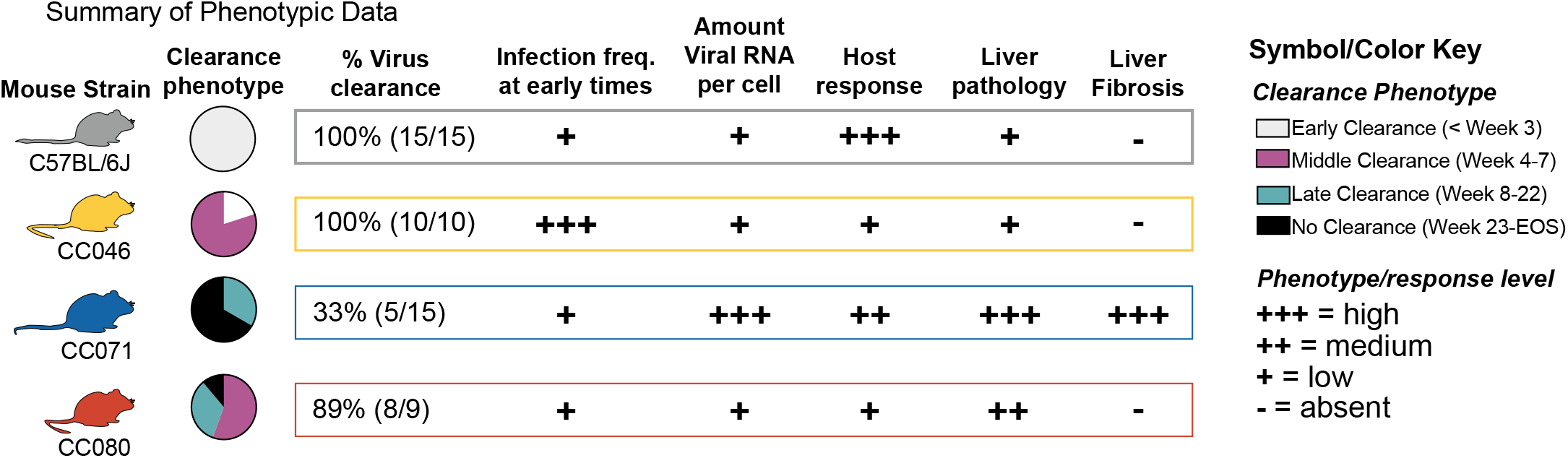
Summary of key phenotypic data per mouse strain. Key phenotypic data per mouse strain is shown to provide an overview of the main pieces of data described in the manuscript.

## Notes

### Competing Interest Statement

The authors have declared no competing interest.

